# Parallel evolution of X chromosome-specific SMC complexes in two nematode lineages

**DOI:** 10.1101/2024.05.21.595224

**Authors:** Avrami Aharonoff, Jun Kim, Aaliyah Washington, Sevinç Ercan

**Affiliations:** Department of Biology, Center for Genomics and Systems Biology, New York University, New York, NY 10003

**Keywords:** dosage compensation, sex chromosomes, X chromosome, nematodes, steinernema, pristionchus, caenorhabditis, oscheius, Hi-C, TAD, condensin, Structural Maintenance of Chromosomes, SMC complexes, evolution, parallel evolution, transcription, H4K20me1

## Abstract

Mechanisms of X chromosome dosage compensation have been studied extensively in three model organisms that represent distinct clades. The diversity within each clade as a function of sex chromosome evolution though is largely unknown. Here, we anchor ourselves to the nematode *Caenorhabditis elegans*, where dosage compensation is accomplished by an X chromosome specific condensin that belongs to the family of structural maintenance of chromosomes (SMC) complexes. By combining a phylogenetic analyses of the *C. elegan*s dosage compensation complex with a comparative analysis of its epigenetic signatures, such as X-specific topologically associating domains (TADs) and enrichment of H4K20me1, we show that the condensin-mediated mechanism evolved recently in the lineage leading to *Caenorhabditis* following an SMC-4 duplication. Unexpectedly, we found an independent duplication of SMC-4 in *Pristionchus pacificus* along with the presence of X-specific TADs and H4K20me1 enrichment, which suggests that condensin-mediated dosage compensation evolved more than once in nematodes. Differential expression analysis between sexes in several nematode species indicates that dosage compensation itself precedes the evolution of X-specific condensins. In Rhabditina, X-specific condensins may have evolved in the presence of an existing mechanism linked to H4K20 methylation as *Oscheius tipulae* X chromosomes are enriched for H4K20me1 without SMC-4 duplication or TADs. In contrast, *Steinernema hermaphroditum* lacks H4K20me1 enrichment, SMC-4 duplication, and TADs. Together, our results indicate that dosage compensation mechanisms continue to evolve in species with shared X chromosome ancestry, and SMC complexes may have been coopted repeatedly in nematodes, suggesting that the process of evolving chromosome wide gene regulatory mechanisms are constrained.

**Significance statement:** X chromosome dosage compensation mechanisms evolved in response to Y chromosome degeneration during sex chromosome evolution. However, establishment of dosage compensation is not an endpoint. As sex chromosomes change, dosage compensation strategies may have also changed. In this study, we performed phylogenetic, genomic, transcriptomic, and epigenomic analyses in several nematode species surrounding *Caenorhabditis elegans* and found that the condensin mediated dosage compensation mechanism in *C. elegans* is surprisingly new, and evolved in the presence of an existing mechanism. Intriguingly, condensin based dosage compensation may have evolved more than once in the nematode lineage, the other time in *Pristionchus*. Together, our work highlights a previously unappreciated diversity of dosage compensation mechanisms within a clade, and suggests constraints in evolving new mechanisms in the presence of existing ones.

## INTRODUCTION

According to the current model of sex chromosome evolution, heteromorphic X and Y chromosomes descend from a homologous, recombining set of autosomes that acquire a sex determining role^1–5^. Suppression of recombination between these newly formed sex chromosomes leads to the degeneration of the Y chromosome and almost complete hemizygosity of the male X chromosome^1,6,7^. Maintenance of the ancestral gene dose, however, is particularly important for dosage sensitive gene networks with a mix of autosomal and sex chromosomal genes^8^. In order to maintain ancestral gene dose, mechanisms of sex chromosome specific gene regulation are selected for. These genetic and epigenetic mechanisms compensating for the imbalance in gene dose between males and females are collectively referred to as “dosage compensation”^3^.

Sex chromosomes have co-evolved with dosage compensation many times. Therefore, clades defined by a shared ancestral X chromosome can also be defined by their dosage compensation mechanism. For example, mammals, flies, and nematodes all use different strategies. In *Mus musculus* (mouse), a long noncoding RNA, *Xist*, initiates several heterochromatic pathways to randomly inactivate one of the X chromosomes in females (XX)^9–11^. In *Drosophila melanogaster* (fly), a histone acetylating ribonucleoprotein complex, *male specific lethal* (MSL) activates transcription by about two-fold on the single X chromosome in males (XY)^12,13^. Finally, in the nematode *Caenorhabditis elegans*, the *dosage compensation complex* (DCC) is driven by an X-specific Structural Maintenance of Chromosome (SMC) complex that represses transcription by about two-fold in hermaphrodites (XX)^14–18^ (**Figure 1A**).

**Figure 1.**
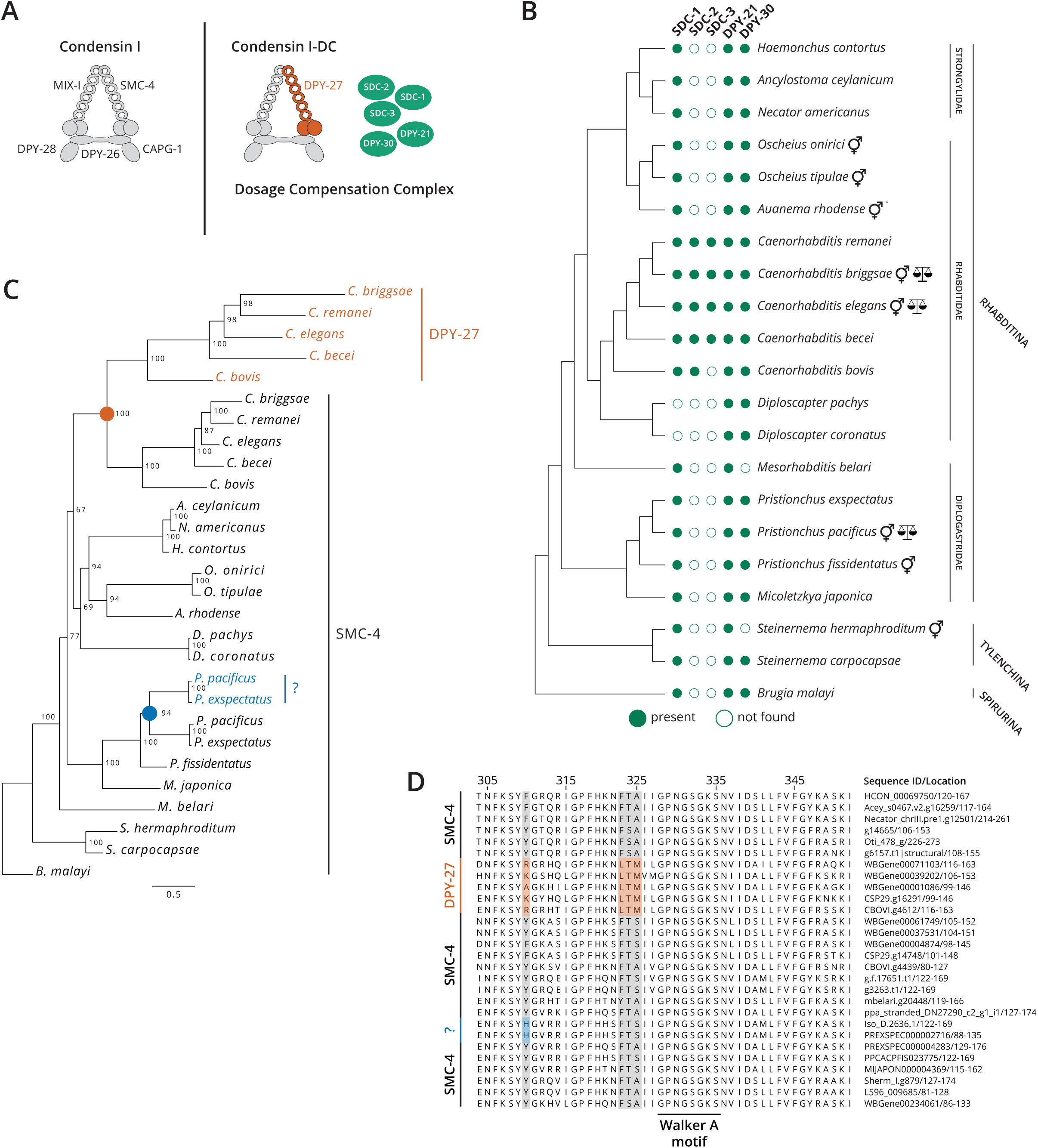
Independent duplications of SMC-4 in *Caenorhabditis* and *Pristionchus* generated novel SMC-4 proteins. (A) Condensin I-DC and five non-condensin proteins (green) make the *C. elegans* dosage compensation complex (DCC). Condensin I-DC shares all but one subunit, DPY-27, with the canonical condensin I. (B) Proteomes with BUSCO scores > 85% were run in OrthoFinder and grouped as orthogroups. SDC-2 and SDC-3 only appear in *Caenorhabditis*, whereas SDC-1, DPY-21, and DPY-30 are conserved among these three clades. *trioecious. (C) OrthoFinder results were used to generate a maximum-likelihood phylogeny of SMC-4. DPY-27 is an SMC-4 paralog that arose in the lineage leading to *Caenorhabditis* (orange circle). An independent duplication of SMC-4 occurred in *Pristionchus* (blue circle). For simplicity, species names were substituted for gene names. Gene names can be found in Table 2. Numbers indicate bootstrap values. Scale indicates branch length. (D) Multiple sequence alignment of the N-terminal ATPase domain highlights conserved and distinct amino acid sequences in DPY-27 and SMC-4. Within this region, example amino acid changes that support the duplication and divergence of DPY-27 in *Caenorhabditis* are shown in orange, and changes that support the independent duplication of SMC-4 in *Caenorhabditis* and *Pristionchus* are shown in blue.

The descent of heteromorphic X and Y chromosomes is *not* an end point. Chromosomal duplications, deletions, fusions, and translocations continue to change sex chromosomes^19–23^. Therefore, although initial dosage compensation mechanisms evolve in response to Y chromosome degradation, they continue to evolve in response to X chromosome evolution. For example, in two species of *Drosophila*, a translocation of autosomal sequences to the X chromosome selected for DNA sequence motifs capable of recruiting the MSL complex to the neo-X region^24–26^. While the motifs required for MSL recruitment remained similar in the *Drosophila* system, in two *Caenorhabditis* species, *C. elegans* and *C. briggsae* (estimated to have diverged from *C. elegans* 15-30 million years ago), the DNA sequence motifs that recruit the DCC have functionally diverged^27^.

The studies performed in *Drosophila* used species in which MSL complex subunits were readily identified by sequence homology^24–26^. Similarly, subunits of the *C. elegans* DCC were identified by sequence homology in *C. briggsae*^27^. Studying the mechanism of dosage compensation in more distantly related species, however, is difficult due to the rapid evolution of proteins involved in sex determination and dosage compensation^2,21^. Furthermore, until recently, chromosome scale assemblies in non-model organisms were few and far between. Therefore, the field remains largely ignorant to the diversity in dosage compensation mechanisms in species that have diverged significantly, but share an X chromosome ancestor.

Here, we used a combination of phylogenetic and genomic approaches to explore dosage compensation in several species of nematodes with significant differences in X chromosome content. We anchored our work in *C. elegans*, a model organism in which XX and XO animals develop as hermaphrodites and males, respectively^15^. The core of the *C. elegans* DCC is a condensin complex belonging to the SMC family^17,28–31^. Most animals harbor two canonical condensins (I and II) that mediate DNA loops to compact chromosomes, and are essential for chromosome segregation in mitosis and meiosis^29^. Condensins I and II share a heterodimer of SMC-2 and SMC-4 subunits, which interact with a different set of three chromosome-associated proteins (CAP)^32^. In *C. elegans*, the third condensin (I^DC^) is formed by replacing SMC-4 with DPY-27 in condensin I (**Figure 1A**)^17^. Condensin I^DC^ (hereafter I-DC) interacts with five non-condensin proteins to form the DCC^33^.

The two major epigenetic changes mediated by the DCC on the X chromosomes are (1) the formation of loop-anchored topologically associating domains (TADs) and (2) the enrichment of H4K20me1 in females and hermaphrodites^34–37^. To explore dosage compensation across distantly related nematodes, we first generated a phylogeny of SMC-4 and DPY-27 in Rhabditina, Tylenchina, and Spirurina. We then looked for condensin-mediated epigenetic phenotypes by performing Hi-C and ChIP-seq in several species. Based on the presence of two SMC-4 paralogs, X-specific loop anchored TADs, and X enrichment of H4K20me1, we conclude that condensin-mediated dosage compensation arose more than once, the other time being in *Pristionchus.* In *Oscheius*, we found evidence for dosage compensation and upregulation of H4K20me1, but not for TADs, and in *Steinernema*, we found an absence of dosage compensation, TADs, and H4K20me1. We surmise that dosage compensation is ever changing in nematodes, and that the mechanism in the common ancestor of *Oscheius, Caenorhabditis*, and *Pristionchus* may be through the well conserved histone H4 Lysine 20 demethylase DPY-21. We propose that the X-specific condensin arose in the lineage leading to *Caenorhabditis* in the presence of this mechanism. Our findings highlight the continued and constrained evolution of dosage compensation mechanisms, and a previously underappreciated diversity of strategies in species with a shared X chromosome ancestry.

## MATERIALS AND METHODS

### Phylogenetic analysis

Datasets used can be found in **supplemental file 1**. To assess annotation quality, BUSCO scores were called in protein mode against the nematoda lineage. We used version 3.0.2 (OBD9 dataset) for the diplogastrids, and version 5.3.0 (ODB10 dataset) for the rest^38–41^. We called orthologs using Orthofinder (version 2.5.4) by first extracting the longest transcript per gene with the provided primary_transcripts.py script, and then running Orthofinder with the default parameters^42,43^. We removed orthologs that were less than half the size of SMC-4 in *C. elegans*. These sequences represent either partial duplications or duplications followed by deletions, and are unlikely to have the function of SMC-4 preserved. Furthermore, the resulting gaps in the alignment would affect downstream analysis. We aligned orthologs using MAFFT (version 7.475) with the following parameters: --local pair --maxiterate 1000^44^. We generated a maximum-likelihood gene tree using IQ-Tree (version 2.2.0) on the aligned sequences with the following parameters: --seqtype AA -m MFP -B 1000^45–47^. The trees were rooted and edited in FigTree (version 1.4.4) and TreeViewer^48^.

### Nigon painting

Datasets used can be found in **supplemental file 1**. BUSCO (version 5.3.0) was run on genome mode against the nematoda lineage in the ODB10 dataset^38–40^. The full_table.tsv file was used to paint chromosomes online at https://pgonzale60.shinyapps.io/vis_alg/ following the instructions in the github repository https://github.com/pgonzale60/vis_ALG^49^.

### Strains and growth conditions

We used the following strains for *Oscheius tipulae*, *Pristionchus pacificus*, and *Steinernema hermaphroditum*, respectively: CEW1, PS312, and PS9179. *O. tipulae* and *P. pacificus* were maintained on 6 cm NGM plates (3 g NaCl, 2.5 g peptone, 20 g agar, 1 mM calcium chloride, 5 mg/L cholesterol, 1 mM magnesium sulfate, and 25 mM potassium phosphate buffer) seeded with a streptomycin resistant strain of *Escherichia coli*, OP-50. *S. hermaphroditum* was maintained on 10 cm NGM plates seeded with a kanamycin and streptomycin resistant strain of *Xenorhabdus griffiniae*, HGB2587. Plates were cultured at 22, 20, and 25°C, respectively.

### Hi-C

*O. tipulae* and *P. pacificus* were propagated for collection on 10 cm NGM plates supplemented with agarose (10 g agar, 7 g agarose) to prevent burrowing, and seeded with OP-50. Plates were fed as needed with a concentrated strain of *E. coli*, HB101. For *O. tipulae*, mixed stage plates were washed with sterile M9 and a cell spreader to unstick the embryos from the plate. Adults and larvae were killed by nutating in a bleach solution (0.5 M NaOH, 1% sodium hypochlorite) for 2 minutes, washed with M9, and arrested at L1 by nutating in M9 for 24 hours. The worms were filtered through a 15 µ nylon mesh to separate arrested L1 larvae from carcasses and unhatched embryos, and then plated. The worms were cultured for 24 hours, fixed by nutating in 2% formaldehyde (in M9) for 30 minutes, washed with M9, flash frozen in liquid nitrogen, and stored at -80 °C. The extent of germline proliferation after 24 hours was determined by DIC microscopy.

For *P. pacificu*s, mixed stage plates were filtered through a 20 µ nylon mesh to produce a predominantly J2-J3 flow through, which was fixed by nutating in 2% formaldehyde (in M9) for 30 minutes, washed with M9, flash frozen in liquid nitrogen, and stored at -80 °C.

For Hi-C, around 50 µL of pelleted larvae were removed from -80 °C and resuspended in PBS with 1 mM PMSF added fresh. A mortar and pestle (BioSpec Products, catalog # 206) was placed on dry ice and filled with liquid nitrogen. Drops of larvae were pipetted into the mortar, and pulverized into a fine powder while submerged in liquid nitrogen. The powdered larvae was crosslinked as described in the Arima Hi-C+ High Coverage Kit (catalog # A101030). Hi-C (restriction enzymes: DpnII, HinfI, DdeI, and MseI) was then performed as per Arima’s instructions. Library preparation was performed with the KAPA Hyper Prep Kit (KK8502) as per Arima’s instructions. Paired-end (100 bp) sequencing was performed at the Genomics Core at the Center for Systems and Genome Biology, New York University using the Illumina NovaSeq 6000 platform.

### Re-scaffolding and liftover

Hi-C fastq files were mapped to the *P. pacificus* genome (PRJNA12644), and biological replicates were merged using the Arima Genomics mapping pipeline (https://github.com/ArimaGenomics/mapping_pipeline). The output bam file was used to re-scaffold the *P. pacificus* genome with YaHS (version 1.1)^50^. The following parameters were used: -r 1000, 2000, 5000, 10000, 20000, 50000, 100000. The draft genome was curated with Juicebox Assembly Tools (JBAT) in Juicebox (see code availability)^51^. Liftoff (version 1.6.3) was used to liftover the protein coding genes to the updated assembly with the default parameters.^52^

### Hi-C data processing and analysis

Hic files were generated using Juicer (version 1.5.7)^51,53^. Biological replicates were merged using Juicer’s mega.sh script. All output hic files (minimum mapping quality of 30) were converted to cool files using the hicConvertFormat tool in HiCExplorer (version 3.6) with the following parameters: --inputFormat hic --outputFormat cool. Count data was loaded from the cool file using the following parameters: -- inputFormat cool --outputFormat cool --load_raw_values. This raw cool file was then balanced using Cooler (version 0.8.11) with the following parameters: --max-iters 500 --mad-max 5 --ignore-diags 2. The balanced cool file was used in Cooltools (version 0.4.0) to generate the Hi-C matrixes, calculate observed/expected and insulation scores, and compute the P(s) and its derivative (see code availability)^54^. All plots were made with Matplotlib (version 3.4.3)^55^.

### RNA-seq data processing and analysis

We used the nf-core rna-seq pipeline (version 3.12.0) to generate counts and TPM tsv files. The workflow of the pipeline under our parameters was as follows: catenate technical replicates, infer strandedness with Salmon (version 1.10.1), asses the sequencing quality with FastQC (version 0.11.9), remove rRNA reads with sortMeRNA (version 4.3.4), align with STAR (version 2.7.10a), quantify with Salmon (version 1.10.1), and summarize counts and TPM quantification at the gene level with tximport (version 1.16.0). We used the follow flags: -profile singularity, --remove_ribo_rna, --save_non_ribo_reads, --save_reference, --skip_umi_extract, --skip_trimming, --skip_bbsplit_reads, --skip_biotype_qc, --skip_stringie, --skip_deseq2_qc. We ran DESeq2 (version 1.42.0) on the gene counts tsv file in R, and used ggplot2 (version 3.5.0) for all plots^56,57^. We used the gene TPM tsv files to filter out unexpressed genes (mean TPM of replicates > 1 in both sexes).

### ChIP-seq

*O. tipulae* and *P. pacificus* were collected like they were for Hi-C. *S. hermaphroditum* was collected as follows. Thirty infective juveniles (F_0_) were split among 3 10 cm NGM plates seeded with HGB2587, and fed as necessary with a concentrated form of the same strain until the second generation (F_1_) reached adulthood. The adults were moved with sterile water onto 9 10 cm plates in anticipation of overcrowding in the third generation (F_2_). After a couple of days, when the plates were concentrated with third generation larvae, the worms were filtered through a 30 µ filter to produce a larval population without extensive germline proliferation. They were then fixed by nutating in 2% formaldehyde (in PBS) for 30 minutes, washed with PBS, flash frozen in liquid nitrogen, and stored at -80 °C. The extent of germline proliferation was determined by DIC microscopy.

Around 100-150 µL of pelleted larvae were removed from -80 °C and washed in FA Buffer (50 mM HEPES/KOH pH 7.5, 1 mM EDTA, 1% Triton X-100, 0.1 % sodium deoxycholate, 150 mM NaCl, filter sterilized) with 0.1% sarkosyl, 1 mM PMSF, and 1X protease inhibitors (protease inhibitor cocktail set I – Calbiochem, catalog # 539131) added fresh. Larvae were then dounced 30 times in a glass dounce tissue grinder (VWR, catalog number 22877-280) with pestle type B. Sonication was performed in a Biopruptor ® Pico (catalog # B01060010) to obtain fragments between 200 and 800 bp in length (30 seconds on, 30 seconds off, 10-15 cycles). Protein concentration was determined using the Bradford assay (Bio-Rad, catalog # 500-0006). For chromatin immunoprecipitation, 1-2 mg of extract and 3-5 µg of antibody were used (see **supplemental file 1**), and allowed to rotate overnight at 4 °C. nProtein A Sepharose beads (Cytiva, catalog # 17528001) were added and incubation was allowed to continue for 2 hours.

Beads were washed with FA buffer, FA buffer – 1 M NaCl, FA buffer – 500 mM NaCl, TEL buffer (0.25 M LiCl, 1% NP-40, 1% sodium deoxycholate, 1 mM EDTA, 10 mM Tris-HCl, pH 8.0), and TE, respectively. Immunocomplexes were eluted twice with ChIP elution buffer (1% SDS, 250 mM NaCl, 10 mM Tris pH 8.0, 1 mM EDTA) at 65 °C for 15 min at 1400 rpm. The eluate was then treated with 2 µL proteinase K (10 mg/mL) for 1 hour at 50 °C, and reverse crosslinked overnight in a 65 °C waterbath. DNA was purified the next morning with Qiagen MinElute PCR Purification Kit (catalog # 28006), and fragment size was checked by agarose gel electrophoresis or TapeStation. The remaining DNA was stored at -20 °C.

Library preparation was performed as follows. To perform end repair, we used half the ChIP DNA and 10-30 ng of input DNA, T4 DNA Ligase Reaction Buffer (NEB, catalog # B0202S), 0.4 mM dNTPs, 20 U PNK (Fisher, catalog # FEREK0032), 0.7 U Large Klenow Fragment (NEB, catalog # M0210S), and 6 U T4 DNA Polymerase (Fisher, catalog # FEREP0062) (total volume 33 µL). The reaction was incubated at 20 °C for 30 minutes, and was cleaned with Qiagen MinElute PCR Purification Kit (eluted in 21 µL). To perform A-tailing, NEBuffer 2 (catalog # B7002S), 0.2 mM dATP, and 7.5 U of Klenow Fragment (3’ → 5’ exo-) (NEB, catalog # M0212L) were added to the cleaned DNA (total volume of 25 µL), and incubated for an hour in a 37 °C water bath. The reaction was cleaned with Qiagen MinElute PCR Purification Kit (eluted in 17 µL). To perform adapter ligation, Quick Ligation Buffer, Illumina Truseq Adapters (see **supplemental file 1**), and 2 µL of Quick Ligase (NEB, catalog # M2200L) were added to cleaned DNA (total volume of 40 µL), and incubated for 20 minutes at 23 °C. The reaction was size selected and cleaned with Ampure XP beads (Beckman Coulter, catalog # A63881). The eluate was amplified by PCR with Phusion HF Reaction Buffer, 0.5 µM TruSeq forward and reverse primers (see **supplemental file 1**), 0.2 mM dNTP, and 1 U Phusion High-Fidelity DNA Polymerase (Fisher, catalog F-530L) for a total reaction volume of 50 µL under the following parameters: 98 °C for 1 min, 98 °C for 30 seconds, 60 °C for 30 seconds, 72 °C for 30 seconds, repeat steps 2-4 14 times, 72 °C for 5 min, 8 °C forever. 10 µL of 3 M sodium acetate (pH 5.2) was added to reaction, and the reaction was cleaned with Qiagen MinElute PCR Purification Kit, (eluted in 11 µL). The eluate was gel purified (1.5% agarose gel) to obtain fragments between 250-600 bp using the Qiagen MinElute Gel Extraction Kit (catalog # 28606). Library concentration was determined with the KAPA Library Quantification Kit (catalog # KK4824). Single-end (75 bp) sequencing was performed at the Genomics Core at the Center for Systems and Genome Biology, New York University using the Illumina NextSeq 500 platform.

### ChIP-seq data processing and analysis

When necessary, adapters were trimmed with trimmomatic (version 0.39). Reads were aligned with bowtie2 (version 2.4.2) using the default parameters. Sam files were converted to bam files, sorted and indexed using samtools (version 1.11). Bam files were used to generate bigwig files with the bamCoverage tool in deeptools (version 3.5.0) with the following parameters: --binsize 10 --minMappingQuality 20 --extendReads 200 --ignoreDuplicates --normalizeUsing CPM --exactScaling --ignoreForNormalization chrM. Enrichment was normalized to input with the bigwigCompare tool. Bigwig files were merged with the bigWigMerge tool in deeptools. Genome browser tracks were made with the hicPlotTADs tool in HiCExplorer (version 3.6). Enrichment around transcription start sites (TSS) and gene bodies were computed for each replicate and on the merged bigwig files with the multiBigWigSummary tool in deeptools. TSS sites were defined as 250 bp upstream and downstream of the first base pair. The outputs were z-scored and plotted in R with ggplot2 (version 3.5.0)^57^.

### Statistics

Wilcoxon rank sum tests were run in R using the wilcox.test function. Permutation tests were also run in R, and as follows. Autosomes were collapsed, and the observed difference in mean loop size between the autosomes (A) and the X chromosomes (X) was calculated as our test statistic. Mean loop size was defined as the separation in base pairs at the local maxima of the log derivative of the P(s)^58–60^. Permutations were run (n=10000) by extracting slope values, and randomly sampling a chromosome (A or X) without replacement. For each permutation, the mean loop size of A and X was selected, and the difference was calculated. A distribution of differences was plotted with ggplot2 (version 3.5.0). We calculated the p-value as a proportion of permuted test statistics that were more extreme than the observed test statistic.

### Code availability

All scripts run in this study were deposited to the following Github repository: https://github.com/ercanlab/2024_Aharonoff_et_al.git.

### Data availability

The datasets used and generated in this paper are provided in **supplemental file 1**. All data generated in this study were deposited to the Gene Expression Omnibus database under series numbers GSE267962 and GSE267963. The re-scaffolded *P. pacificus* genome and lifted annotation of protein coding genes can be found at https://github.com/ercanlab/2024_Aharonoff_et_al.git

## RESULTS

### Ortholog analysis reveals variable levels of conservation among DCC subunits

Nematodes represent a large number of species with distinct genomes. We used a recent phylogenomic analysis to choose a set of species from Rhabditina (clade V) and Tylenchina (clade IV)^61^ (**Figure 1B**). All species selected had BUSCO scores of at least 85% with the exception of *S. carpocapsae*, which fell at 76% (**Table 1**). In all, we sampled from the Strongylidae, Rhabditidae, Dipoglastridae, and Steinernematidae families, which display significant divergence in sex chromosome content (**Figure S1**)^49,62,63^. Our outgroup was *Brugia malayi* (Spirurina, clade III). To determine the conservation of the DCC, we searched for the orthologs with Orthofinder, which uses a reciprocal best blast hits (RBBH) approach.

**Table 1.** Species sampled for phylogenetic analysis of dosage compensation proteins.

| Species | Clade | Family | BUSCO (S/D/F/M) |
| --- | --- | --- | --- |
| *Haemonchus contortus* | Rhabditina | Strongylidae | 85/11/1/3 |
| *Ancylostoma ceylanicum* | Rhabditina | Strongylidae | 86/9/2/3 |
| *Necator americanus* | Rhabditina | Strongylidae | 46/51/1/2 |
| *Oscheius onirici* | Rhabditina | Rhabditidae | 89/3/2/6 |
| *Oscheius tipulae* | Rhabditina | Rhabditidae | 88/3/2/8 |
| *Auanema rhodense* | Rhabditina | Rhabditidae | 90/1/1/8 |
| *Caenorhabditis remanei* | Rhabditina | Rhabditidae | 95/4/1/0 |
| *Caenorhabditis briggsae* | Rhabditina | Rhabditidae | 80/19/0/0 |
| *Caenorhabditis elegans* | Rhabditina | Rhabditidae | 66/34/0/0 |
| *Caenorhabditis becei* | Rhabditina | Rhabditidae | 89/9/1/2 |
| *Caenorhabditis bovis* | Rhabditina | Rhabditidae | 77/17/1/6 |
| *Diploscapter pachys* | Rhabditina | Rhabditidae | 89/2/2/8 |
| *Diploscapter coronatus* | Rhabditina | Rhabditidae | 89/2/2/8 |
| *Mesorhabditis belari* | Rhabditina | Rhabditidae | 72/13/2/13 |
| *Pristionchus exspectatus* | Rhabditina | Diplogastridae | 90/3/4/4 |
| *Pristionchus pacificus* | Rhabditina | Diplogastridae | 96/2/2/0 |
| *Pristionchus fissidentatus* | Rhabditina | Diplogastridae | 89/4/5/2 |
| *Micoletzky japonica* | Rhabditina | Diplogastridae | 88/2/5/5 |
| *Steinernema hermaphroditum* | Tylenchina | Steinernematidae | 43/45/1/11 |
| *Steinernema carpocapsae* | Tylenchina | Steinernematidae | 58/18/4/21 |
| *Brugia malayi* | Spirurina | Onchocercidae | 68/31/0/1 |

*S*, Complete and single copy. *D*, Complete and duplicated. *F*, Fragmented. *M*, Missing. Completeness is measured by summing *S* and *D*.

The DCC subunits SDC-1, SDC-2, and SDC-3 function in both sex determination and dosage compensation^64^. It is well documented that sex determination mechanisms evolve with relative rapidity and are highly flexible to change^21,65^. SDC-2 is expressed specifically in XX embryos, and initiates hermaphrodite development and X chromosome dosage compensation^66^. SDC-3 physically interacts with SDC-2, helping repress the male determination gene *her-1* and recruit the DCC to the X chromosomes^64^. Consistent with their role as upstream regulators of sex determination, we only find orthologs of SDC-2 and SDC-3 in *Caenorhabditis* (**Figure 1B**).

Unlike SDC-2 and SDC-3, SDC-1 is not an essential gene, and is not required for DCC binding to the X chromosomes^67^, but contributes to *her-1* repression^64,68–70^. We found SDC-1 orthologs in all except for *Diploscapter* (**Figure 1B**). In *C. bovis, C. becei, C. elegans, C. briggsae, C. remanei, M. japonica, S. hermaphroditum*, and *S. carpocapsae*, multiple orthologs were identified (see **supplemental file 1**). SDC-1 is a predicted DNA binding transcription factor (TF) with seven zinc fingers^71^. It is possible that the conservation and expansion of SDC-1 is related to its TF function in hermaphrodite differentiation.

The DPY-21 subunit of the DCC contains clear orthologs across all species analyzed, and DPY-30 in all but two (**Figure 1B**). DPY-21 is a conserved histone demethylase that is expressed in both the germline and soma with functions outside dosage compensation^72^.

DPY-30 is a subunit of both the DCC and COMPASS, a chromatin modifying complex that is conserved across species^73–75^. Within COMPASS, DPY-30 interacts with ASH-2, and is required for trimethylation of H3K4, which is associated with active gene promoters^73,76,77^. The non-dosage compensation roles of DPY-21 and DPY-30 in chromatin regulation may contribute to their deeper conservation.

### Phylogenetic analysis of SMC-4 reveals two independent duplications in *Caenorhabditis* and *Pristionchus*

In condensin I-DC, DPY-27 replaces SMC-4, one of the two ATPase subunits of the canonical condensin I. To determine whether DPY-27 is a paralog of SMC-4, and to uncover the timing of the proposed duplication, we identified SMC-4 homologs from the OrthoFinder results and produced a maximum-likelihood tree of the sequences (**Figure 1C**, **Table 2**). Our tree suggests that DPY-27 is an SMC-4 paralog and is at least as old as the common ancestor of *C. elegans* and *C. bovis*, but younger than the common ancestor shared between *Caenorhabditis* and *Diploscapter* (**Figure 1C**).

**Table 2.**
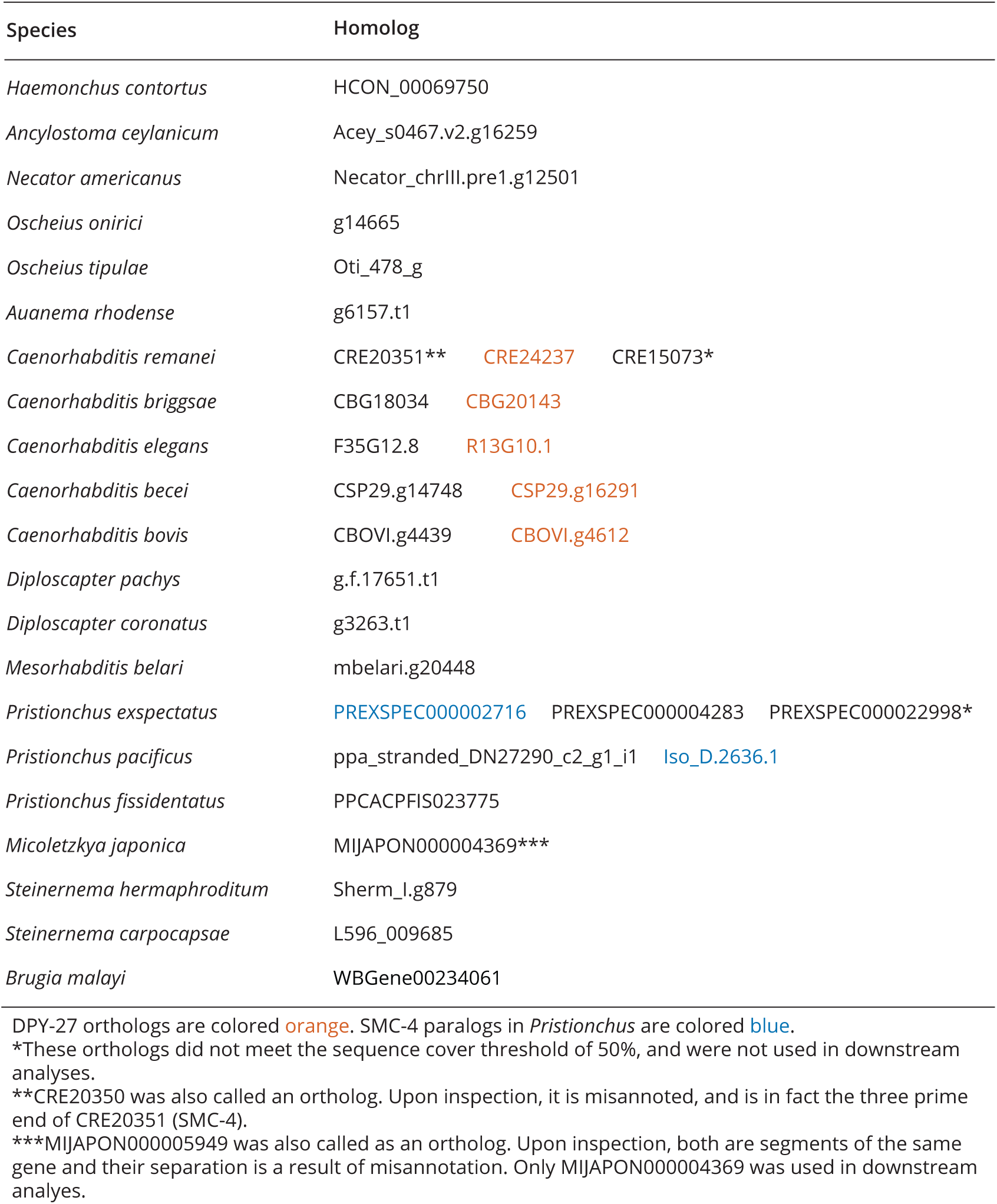
Homologs of *C. elegans* SMC-4.

| Species | Homolog |  |  |
| --- | --- | --- | --- |
| *Haemonchus contortus* | HCON_00069750 |  |  |
| *Ancylostoma ceylanicum* | Acey_s0467.v2.g16259 |  |  |
| *Necator americanus* | Necator_chrIII.pre1.g12501 |  |  |
| *Oscheius onirici* | g14665 |  |  |
| *Oscheius tipulae* | Oti_478_g |  |  |
| *Auanema rhodense* | g6157.t1 |  |  |
| *Caenorhabditis remanei* | CRE20351** | CRE24237 | CRE15073* |
| *Caenorhabditis briggsae* | CBG18034 | CBG20143 |  |
| *Caenorhabditis elegans* | F35G12.8 | R13G10.1 |  |
| *Caenorhabditis becei* | CSP29.g14748 | CSP29.g16291 |  |
| *Caenorhabditis bovis* | CBOVI.g4439 | CBOVI.g4612 |  |
| *Diploscapter pachys* | g.f.17651.t1 |  |  |
| *Diploscapter coronatus* | g3263.t1 |  |  |
| *Mesorhabditis belari* | mbelari.g20448 |  |  |
| *Pristionchus exspectatus* | PREXSPEC000002716 | PREXSPEC000004283 | PREXSPEC000022998* |
| *Pristionchus pacificus* | ppa_stranded_DN27290_c2_g1_i1 | Iso_D.2636.1 |  |
| *Pristionchus fissidentatus* | PPCACPFIS023775 |  |  |
| *Micoletzkyia japonica* | MIJAPON000004369*** |  |  |
| *Steinernema hermaphroditum* | Sherm_I.g879 |  |  |
| *Steinernema carpocapsae* | L596_009685 |  |  |
| *Brugia malayi* | WBGene00234061 |  |  |

DPY-27 orthologs are colored orange. SMC-4 paralogs in *Pristionchus* are colored blue.
\*These orthologs did not meet the sequence cover threshold of 50%, and were not used in downstream analyses.
\*\*CRE20350 was also called an ortholog. Upon inspection, it is misannoted, and is in fact the three prime end of CRE20351 (SMC-4).
\*\*\*MIJAPON000005949 was also called as an ortholog. Upon inspection, both are segments of the same gene and their separation is a result of misannotation. Only MIJAPON000004369 was used in downstream analyses.

Interestingly, we observed an independent duplication of SMC-4 in the *Pristionchus* lineage. To examine the paralogs, we performed a multiple sequence alignment of the N and C terminal ATPase domains, as well as regions that were previously found to be conserved and essential for SMC-4 ATPase function^78^. Similar to other SMC subunits, the ATPase activity of DPY-27 is essential for its function^79^. *C. elegans* DPY-27 has the conserved Walker A motif on the N terminus^28^, along with site specific changes to the ancestral SMC-4 sequence (**Figure 1D, Figure S2**). In *Pristionchus pacificus* and *exspectatus*, the Walker A motif is also conserved in both paralogs. Furthermore, we observe site specific changes independent of *Caenorhabditis* upstream of the Walker A motif (**Figure 1D**). We noticed that one group of paralogs in *P. pacificus* and *P. exspectatus* contains conserved changes to the ancestral SMC-4 sequence, suggesting that Iso_D.2636.1 and PREXSPEC000002716 diverged for a new function.

### Hi-C analysis reveals X-specific topologically associating domains in *P. pacificus*

In *C. elegans*, condensin I-DC-mediated DNA loop extrusion increases 3D contacts, and forms loop-anchored TADs specifically on the X chromosomes^36,37^. A small number of cis-regulatory elements that recruit the DCC to the X chromosomes (recruitment elements on the X, *rex*) form the boundaries between TADs, and contact each other over hundreds of kilobase distances, forming *rex*-*rex* loops^36,37^ (**Figure 2A**).

**Figure 2.**
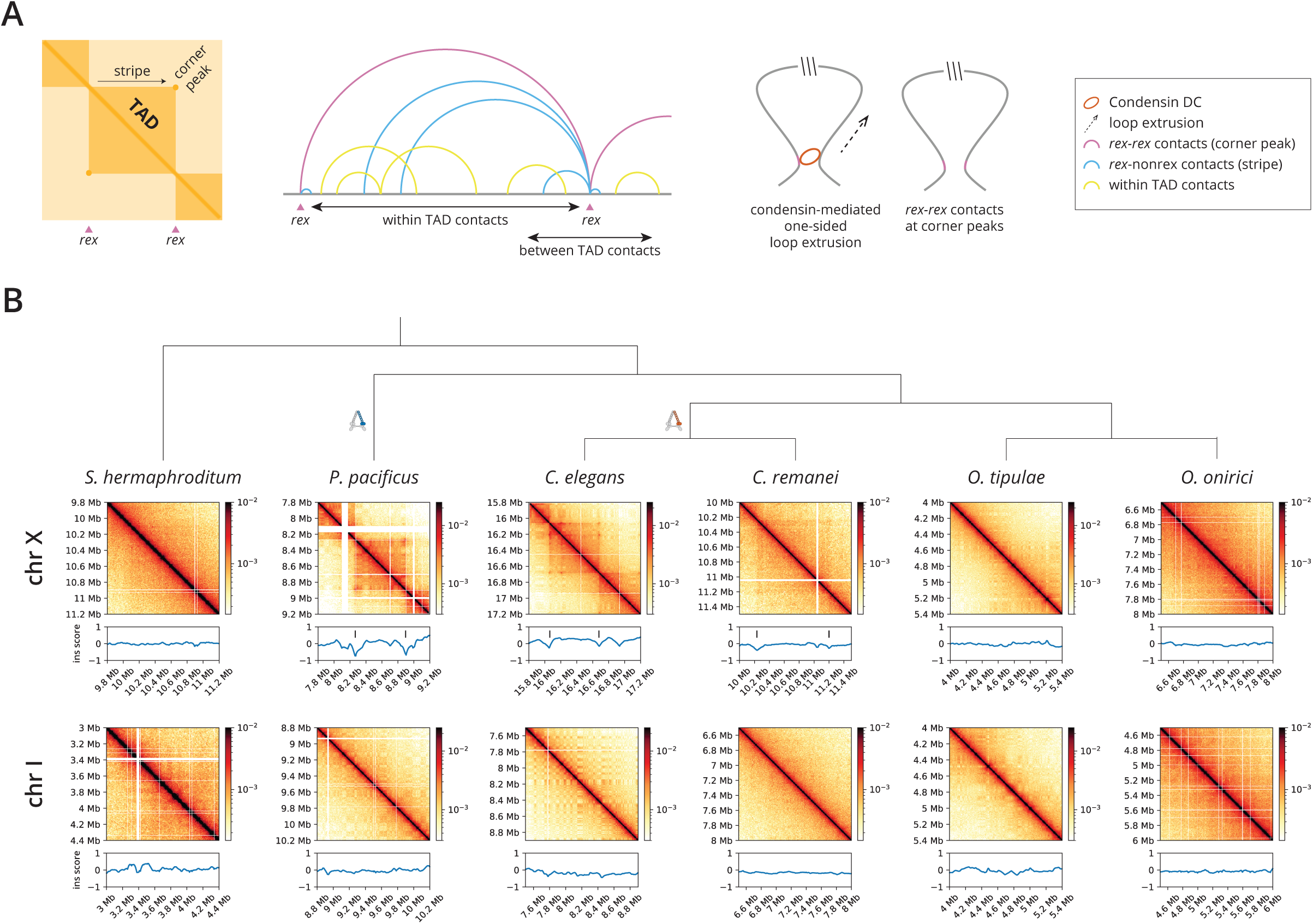
X chromosome specific topologically associating domains are present in *P. pacificus*, but are absent in *S. hermaphroditum*, *O. tipulae*, *and O. onirici*. (A) Schematic diagram of loop-anchored topologically associating domains (TADs) on the *C. elegans* X chromosome. DNA contacts are higher within TADs than between. *C. elegans* TAD boundaries are formed by strong *rex* sites, reminiscent of mammalian CTCF binding elements, and are proposed to function by loading and blocking one-sided DNA loop extrusion of condensin I-DC. (B) Hi-C matrices of representative X and autosome windows at 5 kb resolution with insulation scores below (150 kb window). Empty bins result in insulation score dips that can be tracked by lack of data in the Hi-C matrix (white lines). *P. pacificus* X chromosomes display loop-anchored TADs phenotypic of *C. elegans* dosage compensation along with dips in insulation score at TAD boundaries (tick marks). *S. hermaphroditum*, *O. tipulae* and *O. onirici* do not display X chromosome specific TADs.

To independently validate the results of our phylogenetic analysis, we compared the Hi-C features of the X chromosomes to that of autosomes in six of our sampled species (**Figure 2A**). We generated new data in *P. pacificus* and *O. tipulae* and obtained previously published Hi-C data in *S. hermaphroditum*^80^, *C. elegans*^36^, *C. remanei*^81^ and *O. onirici* (Wellcome Sanger Institute). In each, the X chromosomes are composed of a mosaic of ancestral segments (**Figure S1**)^49,62,63^. Dosage compensation by condensin I-DC is found in the somatic cells of hermaphrodites and females. Therefore, we performed our Hi-C in early stage larvae, before germline cells begin to proliferate (see materials and methods).

While analyzing our Hi-C data mapped to the *P. pacificus* genome, we noticed several misjoins and inversions on chromosomes V and X (**Figure S3**). There were a total of 41 small, unplaced contigs (median length=43,467 bp), many of which displayed contacts with the X chromosome. With our HiC data, we were able to fix the misjoins and inversions on chromosomes V and X, and placed pbcontig517 on the left arm of the X chromosome (**Figure S3**). We also lifted over the annotations of protein coding genes. We re-mapped our *P. pacificus* Hi-C reads to the updated assembly.

In *C. elegans*, condensin I-DC forms loop-anchored TADs specifically on the hermaphrodite X chromosome (**Figure 2B, Figures S4**). In *C. remanei*, all DCC subunits have orthologs, and TAD structures are observed specifically on the X chromosomes (**Figure 2B**). The less defined TAD features are expected because the *C. remanei* Hi-C data was obtained from mixed sex and mixed stage worms, diluting the hermaphrodite and soma-specific features of dosage compensation with male and germ cells.

Strikingly, *P. pacificus* also displayed loop anchored TADs on the X chromosome (**Figure 2B**). While *P. pacificus* has fewer TADs compared to *C. elegans* (**Figure S5**), its X chromosome is less-assembled, which may be affecting their detection. Loop-anchored TADs were absent on the autosomes of all species (**Figure 2B**). Therefore, in *C. elegans*, *C. remanei* and *P. pacificus*, loop-anchored TADs are specific to the X chromosomes.

### Distance decay analysis supports the presence of an X-specific loop extruding factor in P. pacificus

The loop extrusion activity of condensin I-DC on the X chromosomes is also detected by analyzing the decay of DNA contacts as a measure of increased distance, P(s)^36,82^ (**Figure 3, top panel**). Previously, we showed that condensin I-DC mediated DNA loops produce a characteristic shoulder on the distance decay of 3D contacts from the X chromosome compared to the autosomes^36,58,82,83^. We performed the same analysis on our *P. pacificus* Hi-C data, and found that the X chromosome shows the characteristic shoulder not found on the autosomes (**Figure 3, Figure S6**).

**Figure 3.**
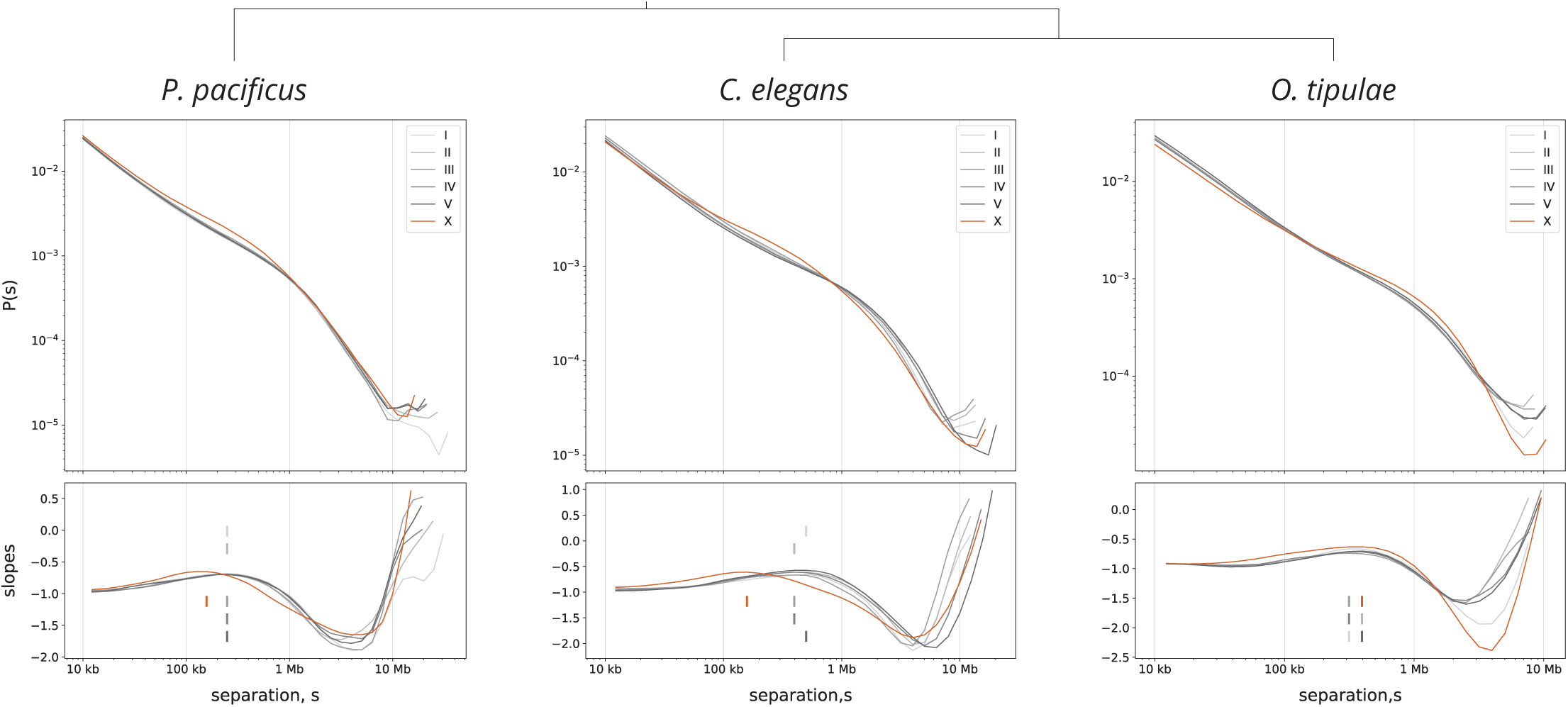
Distance decay curve in *P. pacificus* supports the presence of an X-specific loop extruder similar to the *C. elegans* dosage compensation condensin. Distance decay curves in each species show the contact probability, P(s), as a function of separation, s, for each chromosome at 5 kb resolution (top panel). Mean loop sizes for each chromosome were computed by taking the local maxima of the slope (bottom panel, tick marks). Similar to *C. elegans*, the *P. pacificus* hermaphrodite X chromosomes displays a local maxima shifted to the left (smaller loop size) when compared to the autosomes, indicative of an X chromosome specific loop extruding factor. *O. tipulae* does not show a difference in mean loop size between the X chromosome and autosomes, consistent with a lack of X-specific loop extruding activity.

Modeling of loop extrusion factors (LEFs) suggests that the local maxima of the log-derivative of the P(s) corresponds to the mean loop size, which is decreased by increasing the number of LEFs^58–60^. Indeed, the mean loop size of the *P. pacificus* X chromosome shows the same shift to the left observed in *C. elegans* (**Figure 3, Figure S6**). To test if the mean loop size was statistically different between the X chromosome and autosomes, we performed a permutation test, which supported the presence of an X-specific loop extruder in *P. pacificus* (**Figure S7**). As our test statistic, we collapsed the autosomes, and measured the difference in mean loop size between the autosomes and X chromosomes. In *P. pacificus* and *C. elegans*, the absolute value of the permuted test statistic was greater than the absolute value of the observed in 0.89% and 0.37% of permutations, respectively. In addition, a phylogenetic analysis of cohesin, condensin, and SMC-5/6 subunits found no other SMC duplications (**Figure S8**). Taken with the presence of loop-anchored TADs (**Figure 2B**) and a diverged SMC-4 variant (**Figure 1D**), our data suggest that *P. pacificus* contains a X chromosome specific condensin complex.

### Hi-C analysis indicates no X-specific loop anchored TADs in *O. tipulae*, *O. onirici* and *S. hermaphroditum*

Unlike *P. pacificus*, *O. tipulae* does not contain paralogs of SMC-4 (**Figure 1C**). Importantly, our Hi-C data in *O. tipulae* did not display loop-anchored TADs on the X chromosome (**Figure 2B, Figure S4**). Consistent with the lack of an X chromosome specific loop extruder in *O. tipulae*, there was no difference between mean loop size on the X and autosomes (**Figure 3, Figure S6**). We also analyzed the publicly available Hi-C dataset for *O. onirici* (Wellcome Sanger Institute) and *S. hermaphroditum*, obtained from mixed stage, hermaphrodite animals^80^. While the mixed stage *and* mixed sex *C. remanei* data shows loop-anchored TADs, we observed no TADs in *O. onirici* and *S. hermaphroditum* (**Figure 2B**). Together with our phylogenetic analysis, our Hi-C results place *Oscheius* and *Steinernema* in the lineages lacking an X specific loop extruder.

### *O. tipulae* X chromosomes are dosage compensated

Using mRNA-seq data from males and hermaphrodites, we previously established that *P. pacificus* and *C. remanei* X chromosomes are dosage compensated^84^. In *O. tipulae*, the absence of an SMC-4 paralog and loop-anchored TADs could either be because this species does not compensate or because it uses an alternate mechanism. To test if *O. tipulae* dosage compensates, we analyzed publicly available RNA-seq data^85^ and applied the same criteria established for assessing dosage compensation in *C. elegans*^84,86^.

Dosage compensation mechanisms control the balance between the X and autosomes, and equalize X chromosomal transcript levels between sexes^18,87^. If there is dosage compensation, the average ratio of mRNA expression between sexes should be similar between the X chromosomes and autosomes. We analyzed mRNA-seq data in *B. malayi*, *O. tipulae*, and *H. contortus* adults, and re-analyzed data from *P. pacificus* young adults and *C. elegans* adults^84,85,88–90^. In each case, the average ratio of gene expression between sexes on the X and autosomes was similar (**Figure 4A, top panel**). The expected difference between X and autosomal expression in the absence of dosage compensation is observed in the larvae of a dosage compensation mutant *C. elegans* strain (*dpy-21(y428)*)^35^ (**Figure 4B**).

**Figure 4.**
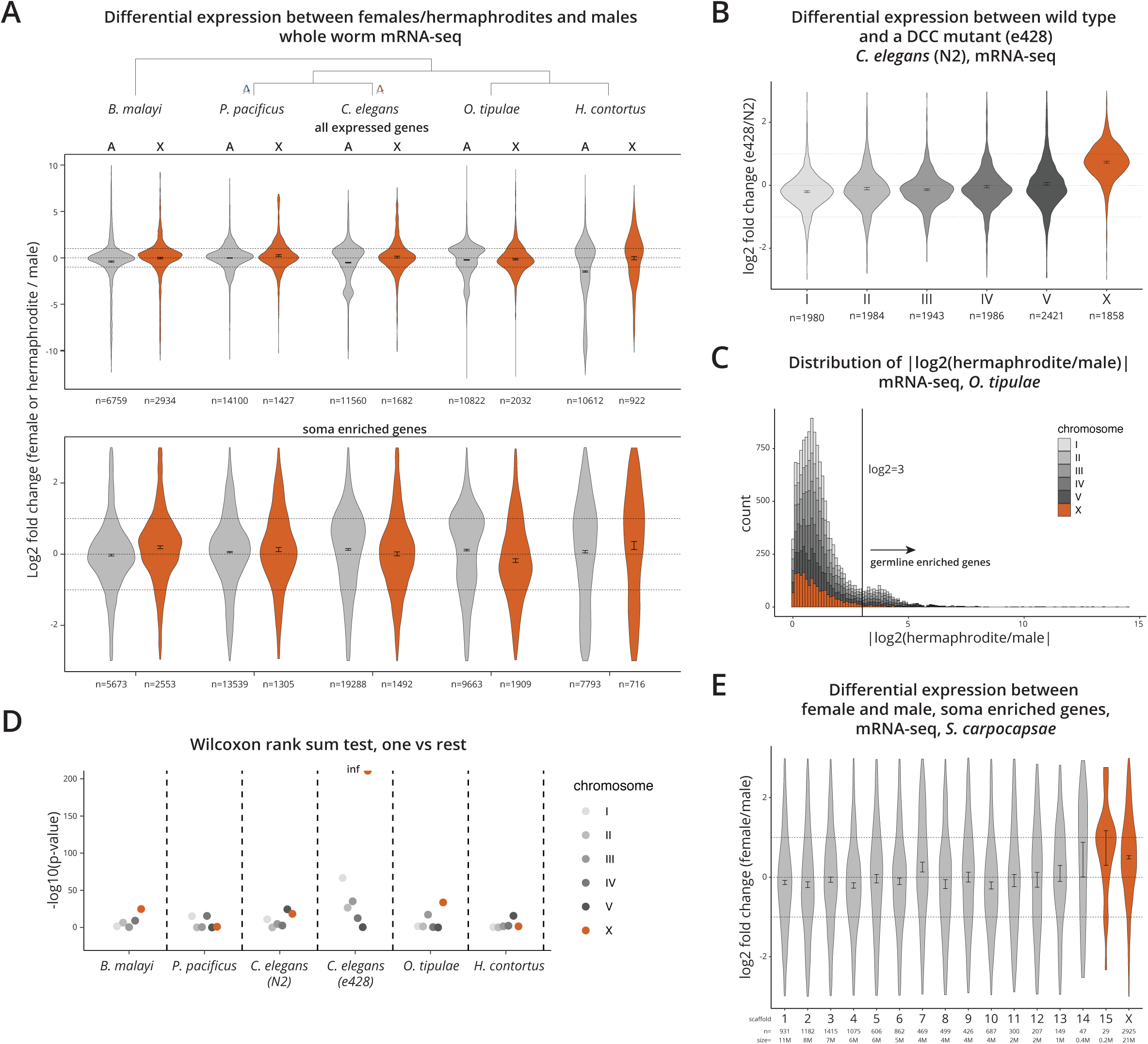
X chromosomes are consistently dosage compensated in multiple nematode lineages. (A) Log2 fold difference in gene expression between females (or hermaphrodites) and males is plotted for expressed genes (mean TPM of replicates > 1 in both sexes) on autosomes and X chromosomes (top panel), and in soma enriched genes (bottom panel). (B) Log2 fold difference between wild type *C. elegans* and a dosage compensation mutant, e428, showing the expected level of expression increases on the X chromosomes in the absence of dosage compensation. (C) Distribution of sex-biased gene expression from adults plotted as |log2(hermaphrodite/male)| in *O. tipulae*. Similar to *C. elegans*, a bimodal distribution of sex-biased gene expression in *O. tipulae* results from germline-enriched genes with higher sex-biased gene expression. (D) -log10(p-value) from Wilcoxon rank sum tests comparing the mean log2 fold difference of a chromosome to the mean of the rest (e.g., mean of X and mean of I-V). p-values in all but the e428 mutant cluster, suggesting no statistical difference between each chromosome in each wild type species. Inf = infinite. (E) Log2 fold difference between females and males in *S. carpocapsae* in soma enriched genes separated by scaffold/chromosome does not support chromosome-wide dosage compensation of the X in this species. Scaffold 15 is colored orange because of its predicted location on the X chromosome^95^.

In *C. elegans*, dosage compensation occurs in the somatic cells, but not in the germ cells, which can make up to two thirds of all cells in adults^86,91^. In addition, genes expressed in the germline are underrepresented on the X chromosomes due to non-sex-specific silencing during meiosis^92,93^. To evaluate dosage compensation in somatic cells, we used our previous strategy of enriching for soma expressed genes based on a cutoff selected by the bimodal distribution of differential expression between sexes^84^.

In *C. elegans*, genes expressed specifically in the germ cells often show greater than 8-fold difference between hermaphrodites (or females) and males ^84^. Thus, using a cutoff of log2 fold change=3 removes most germline-enriched genes^84^. Similar to *C. elegans*, the absolute log2 fold change between sexes shows a bimodal distribution in *O. tipulae* (**Figure 4C**). The level of male-biased gene enrichment on the X chromosome depends on the developmental timing of germline proliferation^84^. We observed varying degrees of male-biased gene depletion in *B. malayi* and *H. contortus*, which may be due to differences in genome evolution or germ cell proliferation (**Figure S9**). Regardless, using the same cutoff across species is a conservative approach to enrich for soma-expressed genes. This resulted in a log2 fold change between sexes to be closer to 0 (**Figure 4A, bottom panel**) and in less variation between autosomes (**Figure S10**).

Each autosome harbors thousands of different genes with varied sex-biased gene expression (**Figure S9, S10**). We used this variability to statistically test for the presence of X chromosome dosage compensation in *O. tipulae*. We employed a “one versus rest” approach, comparing the mean fold difference between sexes on each chromosome to the rest (e.g., I v.s. II, III, IV, V, and X). Similar to *C. elegans* and *P. pacificus*, in *O. tipulae* we found the difference in sex-biased gene expression between the X and autosomes was no more than the difference between an autosome and the rest of the chromosomes (**Figure 4D**). In contrast, in the *C. elegans* dosage compensation mutant (*dpy-21(e428)*), the difference between the X and autosomes is readily apparent. Therefore, X chromosomes are dosage compensated in *O. tipulae*. Together with *B. malayi* and *H. contortus*, these results suggest that dosage compensation precedes condensin I-DC in nematodes, as expected by the model of sex chromosome evolution.

### *Steinernema carpocapsae* X chromosomes are not completely dosage compensated

Similar to *O. tipulae*, *S. hermaphroditum* does not have an SMC-4 duplicate or X-specific loop-anchored TADs. To evaluate if there is dosage compensation in the *Steinernema* lineage, we analyzed published data in *S. carpocapsae*^94^. Although the *S. carpocapsae* genome assembly is not chromosome scale, the X chromosome is assembled, and the remaining four autosomes are split among only 13 contigs^95^. Therefore, we analyzed the mRNA-seq data in female and male young adults. To our surprise, we found that expression in females is higher than males on the X chromosome (log2 fold change > 0), which could not be explained by the natural variation among the autosomes (**Figure 4E, Figure S11**). It is possible that *S. carpocapsae* X chromosomes are not fully dosage-compensated like the other nematode species we analyzed (**Figure 4A**).

### H4K20me1 is enriched on the X chromosomes in both *O. tipulae* and *P. pacificus*

In *C. elegans*, X chromosome repression is mediated by condensin I-DC and DPY-21^34,72,96^. The demethylation of H4K20me2 by DPY-21 results in the enrichment of H4K20me1 on the X chromosomes when compared to autosomes^34,72,96^. The repression of the X chromosomes also results in the depletion of histone modifications associated with active transcription^97^. We wondered if *O. tipulae* and *P. pacificus* show X-specific differences in histone marks associated with dosage compensation in *C. elegans*^34,96,97^. We performed ChIP-seq analysis of H4K20me1, H3K4me3, and as a negative control, IgG in early stage larvae prior to germline proliferation.

We found that similar to *C. elegans*, H4K20me1 is elevated across the entire X chromosome relative to autosomes in both species (**Figure 5A**).

**Figure 5.**
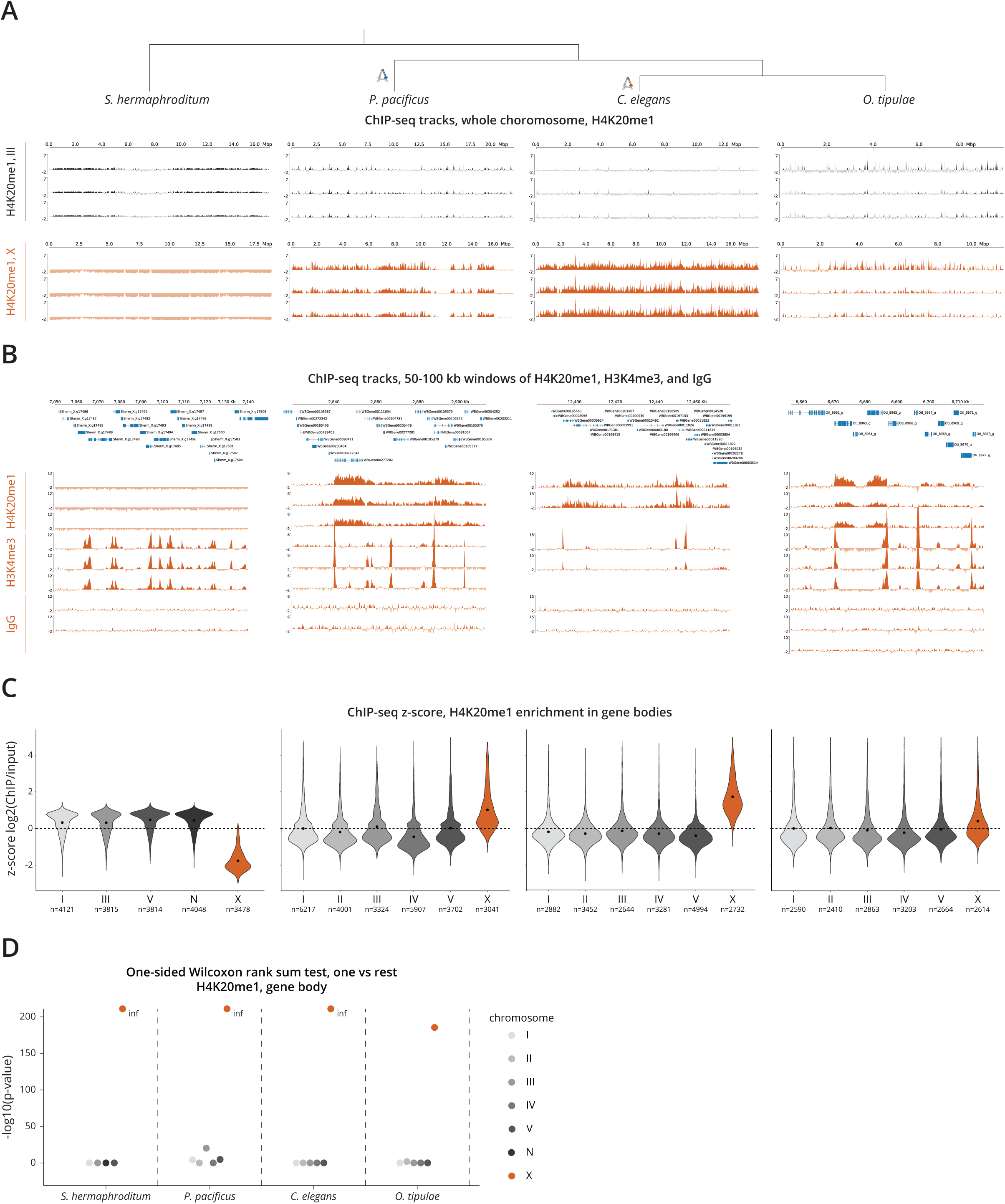
*P. pacificus* and *O. tipulae* X chromosomes are enriched for H4K20me1. (A) H4K20me1 ChIP-seq enrichment tracks (ChIP minus input) are shown for the entire X chromosome and a representative autosome, III in *S. hermaphroditum*, *P. pacificus*, *C. elegans*, and *O. tipulae* hermaphrodite larvae. (B) Representative 50-100 kb windows are shown to highlight the distribution of H4K20me1, H3K4me3, and IgG ChIP-seq tracks (ChIP minus input) relative to gene locations. (C) H4K20me1 ChIP enrichment across gene bodies (z-scored log2(ChIP/input) from transcription start to end site) was plotted for each chromosome. H4K20me1 levels are higher on the X chromosomes of *P. pacificus*, *C. elegans*, and *O. tipulae* when compared to autosomes. In contrast, H4K20me1 is depleted on the X chromosomes of *S. hermaphroditum*. Black dot represents the mean. (D) -log10(p-value) from Wilcoxon rank sum tests comparing the mean log2(ChIP/input) of a chromosome to the mean of the rest (e.g., mean of X and mean of I-V).

H4K20me1 and H3K4me3 ChIP-seq data shown across a representative region highlights the gene-body and promoter enrichment of H4K20me1 and H3K4me3, respectively (**Figure 5B**). This suggests that the functions of these modifications are conserved. Analyzing relative enrichment of H4K20me1 on each chromosome indicated that H4K20me1 is enriched on the X chromosomal gene bodies of each species (**Figure 5C, Figure S12**). We validated that the low enrichment of H4K20me1 on the *O. tipulae* X chromosomes is statistically significant by comparing average gene body ChIP-seq z-scores in each chromosome to the rest (**Figure 5D**).

The enrichment of H4K20me1 in *O. tipulae* is in contrast to the Hi-C data, where loop-anchored TADs are present in *P. pacificus*, but not *O. tipulae*. It is possible that *O. tipulae* has a mechanism of X chromosome repression that employs H4K20 monomethylation without condensin.

### H4K20me1 is depleted on the hermaphrodite X chromosomes of *S. hermaphroditum*

In *Steinernema hermaphroditum*, TADs were absent from the hermaphrodite X chromosome, and in *S. carpocapsae*, we did not find evidence of chromosome-wide dosage compensation. In the possible absence of dosage compensation, we predicted that the *S. hermaphroditum* X chromosome would not be enriched for H4K20me1. ChIP-seq analyses of H4K20me1, H3K4me3, and IgG in *S. hermaphroditum* early stage hermaphrodite larvae showed that the X chromosomes were *depleted* for H4K20me1, and surprisingly, enriched for H3K4me3 (**Figure 5, Figure S12**). This pattern of ChIP-seq enrichment was not a result of the unequal distribution of genes among the chromosomes (**Figure S13**). The absence of H4K20me1 enrichment on the X chromosomes is consistent with both the absence of TADs *and* dosage compensation in *Steinernema*.

## DISCUSSION

In this study, we found dosage compensation mechanisms continue to change within a clade of shared X chromosome ancestry. We also found that the *C. elegans* condensin I-DC evolved recently, in the lineage leading to *Caenorhabditis* and in the presence of an existing mechanism of dosage compensation (**Figure 6**). This mechanism seems to have its basis in the conserved histone demethylase, DPY-21 and may have been present in the common ancestor of Rhabditina. Intriguingly, a condensin based mechanism to regulate X chromosome structure evolved independently in *Pristionchus*, which is indicative of common selective pressures acting on dosage compensation, and a constraint on the evolution of dosage compensation in nematodes.

**Figure 6.**
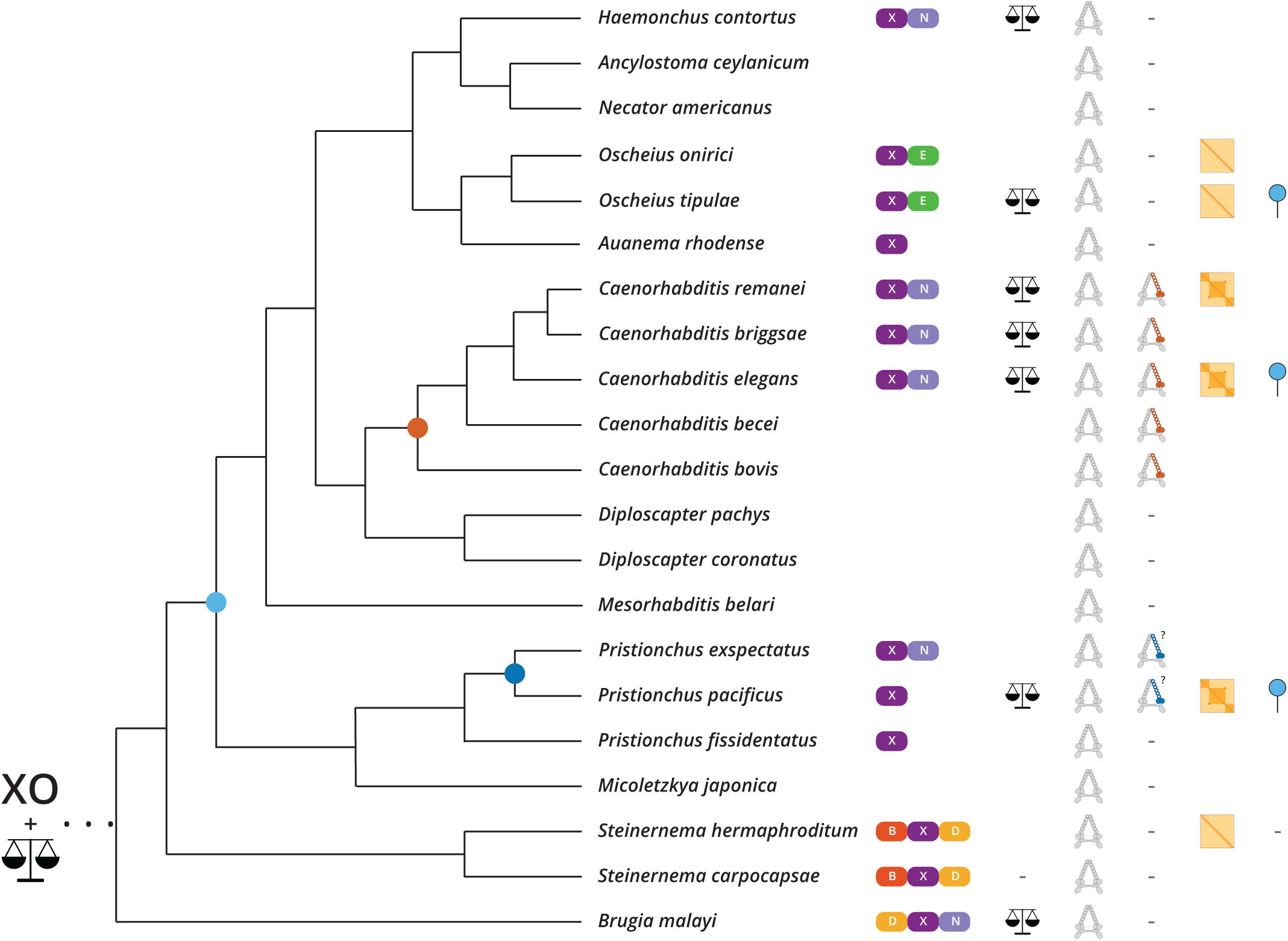
Model for the evolution of dosage compensation in nematodes. Nematodes display significant changes to X chromosome content resulting from autosome to sex chromosome fusions (column 1). Dosage compensation is as old as the common ancestor of Spirurina, Tylenchina, and Rhabditina, which is supported by its presence in *Brugia*, *Pristionchus*, *Caenorhabditis*, *Haemonchus*, and *Oscheius* (column 2). Here, we postulate that it is likely as old as the XO sex determination in nematodes. The canonical SMC-4 is found in all nematodes sampled (column 3). DPY-27, the condensin I-DC specific subunit, is an SMC-4 paralog found only in *Caenorhabditis* (column 4), and the duplication event occurred in the lineage leading to *Caenorhabditis* (orange circle). A second, independent SMC-4 duplication occurred in *Pristionchus* (column 4, dark blue circle). X chromosome specific loop-anchored TADs are a signature of condensin-mediated dosage compensation in *C. elegans*, and are also observed in *P. pacificus* and *C. remanei*, but not in *S. hermaphroditum*, *O. tipulae* and *O. onirici* (column 5). Here, we postulate that X specific condensins in nematodes evolved in parallel in *Caenorhabditis* and *Pristionchus*. The repressive histone mark, H4K20me1 is enriched on the hermaphrodite X chromosomes without SMC-4 duplication or TADs in *O. tipulae*, suggesting it precedes condensins as a mechanism common to Rhabditina (column 6, light blue circle). -, not found; *blank space*, not checked; 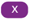, nigon element; 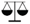, dosage compensation; 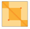, X specific TADS; 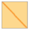 no TADs on X chromosome; 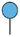, X specific enrichment of H4K20me1.

### XO sex determination and dosage compensation are ancient in nematodes

The degree of Y chromosome degeneration in XY sex determination systems can be a powerful predictor of dosage compensation. For example, in the fish genus *Poecilia*, dosage compensation is found in species with heteromorphic, but not homomorphic sex chromosomes^98^. In nematodes, the complete loss of the Y chromosome is ancestral, although the Y chromosome can reevolve by fusion of the X chromosome with an autosome as observed in *B. malayi*^62^. Analyses of differential gene expression between sexes in *B. malayi*, *O. tipulae*, and *H. contortus* suggest that each of their X chromosomes are dosage compensated (**Figure 4**), as expected from an ancient XO sex determination system in nematodes.

Surprisingly, the *S. carpocapsae* expression data does not support chromosome-wide dosage compensation of its X chromosomes (**Figure 4E**). A parsimonious view of the data along with the age of Y chromosome degeneration in nematodes suggests that dosage compensation was present in the common ancestor of Rhabditina, Tylenchina, and Spirurina, but was lost in the lineage leading to *Steinernema*. This loss may be explained by two significant autosome to sex chromosome fusions in *Steinernema*. The *Steinernema* X chromosome is in fact largely composed of the two autosomal nigon elements B and D, which may not have evolved chromosome-wide dosage compensation following their fusion (**Figure S1**). Furthermore, we postulate that dosage compensation may even be older than the common ancestor of Rhabditina, Tylenchina, and Spirurina, and concurrent with the XO sex determination system (**Figure 6**).

### Condensin-mediated dosage compensation evolved recently in the lineage leading to Caenorhabditis

Our phylogenetic analysis shows that condensin I-DC arose through duplication and divergence of SMC-4 in the lineage leading to *Caenorhabditis* (**Figure 1**). In *C. elegans*, the presence of looped anchored TADs are a molecular phenotype of condensin I-DC activity^36^. While X-specific TADs were found in *C. remanei*, no such TADs were found in *O. tipulae*,*O. onirici*, and *S. hermaphroditum*, which lack an SMC-4 paralog (**Figure 2B**, **Figure 6**). Together, these results suggest that condensin I-DC is a newly evolved mechanism of dosage compensation.

Changes to sex chromosome composition is common in nematodes and may explain the inception of condensin I-DC in *Caenorhabditis*^49,62^. In *Drosophila*, autosome to sex chromosome fusions are followed by degeneration of the non-fused homolog, leaving the neo X-chromosomal genes in one copy^24,26,99^. This dosage problem was solved by evolution of DNA sequence motifs that recruit the existing dosage compensation complex to the neo-X chromosomes. In contrast, different mechanisms of dosage compensation seem to be acting on the neo and ancestral segments of the Z chromosome in the Monarch butterfly, *D. plexippus*^100^. In *D. plexippus*, while the neo-segment of the Z chromosome is upregulated in females (ZW) as in *D. melanogaster*, the ancestral segment of the Z chromosome is downregulated in males (ZZ) as in *C. elegans*^100^. Therefore, new mechanisms of X chromosome regulation may evolve in the presence of an old one.

Our data supports the idea that *C. elegans* condensin I-DC evolved in the presence of an existing mechanism in the nematode clade. It is possible that an X chromosome event was the selective pressure to evolve condensin I-DC, such as the X to N fusion in the lineage leading to *Caenorhabditis* (**Figure S1**)^49,62,63^. Such an event is not obvious in *Pristionchus* where a second X-specific SMC complex has evolved. Furthermore, while the X chromosomes are largely syntenic in *C. elegans* and *C. briggsae*, the DNA sequences that recruit the dosage compensation complex to the X chromosomes functionally diverged between the two species^27^. Therefore, large chromosomal events are not required for the continual evolution of the dosage compensation machinery.

### Parallel evolution of condensin-mediated dosage compensation in *P. pacificus*?

In perhaps the most surprising observation, we identified an independent duplication of SMC-4 in the common ancestor of *P. pacificus* and *P. exspectatus* (**Figure 1**, **Figure 6**). Hi-C analysis in *P. pacificus* revealed X-specific loop-anchored TADs in hermaphrodite larvae (**Figure 2B**). An exploration of the other SMC proteins, namely cohesin, condensin I, condensin II, and SMC-5/6 did not reveal any other duplication events in *Pristionchus* (**Figure S8**). A multiple sequence alignment of the conserved ATPase domains implicated one group of SMC-4 paralogs in dosage compensation (**Figure 1D**). The independent duplication and divergence of SMC-4 towards a possible dosage compensation function in *Caenorhabditis* and *Pristionchus* suggests a common selective pressure in these two lineages. It will be important to determine what constraints lead to the parallel evolution of condensin-based mechanisms of dosage compensation in nematodes.

SMC genes are in fact already known to produce new genes by duplication. In fact, the entire SMC family of genes arose from a common SMC ancestor by several rounds of duplication and neofunctionalization^101–104^. Interestingly, an SMC protein for dosage compensation has evolved in not one, but *two* clades with independent X chromosome ancestry. The first is the nematode condensin I-DC, and the second is SMCHD1 in mammals^105^. Similar to the *C. elegans* DPY-27, SMCHD1 functions in 3D organization and repression of the inactivated X chromosomes in mice and humans^105–109^. It is possible that the chromosome-wide binding capabilities of SMC complexes are well suited to solve the chromosome-wide transcriptional imbalance that necessitates dosage compensation.

### Does a mechanism based on H4K20me1 predispose Rhabditina to evolve a condensin mediated dosage compensation mechanism?

ChIP-seq data in *O. tipulae* shows that H4K20me1 is enriched on the X chromosome, despite the absence of an SMC-4 duplication and X-specific loop-anchored TADS. In *C. elegans*, H4K20me1 is enriched by DPY-21, which is recruited by condensin I-DC to the X chromosomes in mid embryogenesis^34,72,96^. H4K20me1 enhances X chromosome repression for dosage compensation^35,72,110,111^. In *C. elegans*, DPY-21 also functions in the germline without condensin I-DC, where it is required for increasing H4K20me1 on the autosomes, but not the X chromosome^72^.

Our results suggest that H4K20me1 enrichment preceded the SMC-4 duplications in Rhabditina. The alternative interpretation that SMC-mediated dosage compensation preceded H4K20me1 requires that the SMC-4 duplicate be lost in *O. tipulae* without the complete loss of H4K20me1. This is less likely because based on the phylogeny of the species analyzed (**Figure 6**), the SMC-4 duplicate would need to be lost four times and gained once: lost in the common ancestor of the *Diplogastrids* followed by a gain in the common ancestor of *P. pacificus* and *P. exspectatus*, lost in the branches leading to *M. belari* and *Diploscapter*, and lost in the common ancestor of *Auanema*, *Oscheius*, *Necator*, *Ancylostoma*, and *Haemonchus*.

H4K20me1 is a highly conserved histone modification that is enriched on mitotic chromosomes when canonical condensins are active^112^. We speculate that the mechanistic link between H4K20me1 and condensin I contributed to the process of condensin I-DC evolution. Canonical condensin I remains cytoplasmic until the nuclear envelope breaks down during cell division^113,114^. Perhaps the nematode condensin I bound and increased H4K20me1 on the X chromosomes during interphase, while also functioning to compact all chromosomes during mitosis. Condensin I’s dual role in dosage compensation and mitosis may have presented two possibly conflicting constraints. Duplication of the SMC-4 subunit followed by divergence would alleviate one of its constraints, allowing for the separate evolution of two condensin I variants.

Interestingly, an additional SMC may have evolved in *Caenorhabditis* to further mediate the potential interaction between condensin I and I-DC. Mass spectrometry analysis of proteins that interact with DPY-27 found a small SMC-like protein 1 (SMCL-1) in *C. elegans*^115^. SMCL-1 lacks the SMC hinge and coils, contains a non-functional ATPase domain, and negatively regulates condensin I and I-DC^115^. Future studies should address if parallel evolution of the SMC-4 paralog in *P. pacificus* also necessitated the emergence of a negative regulator of condensin I.

### Concluding remarks

While the field has been able to assess whether X chromosomes are dosage compensated or not by using differential expression analysis, the mechanisms of dosage compensation have been traditionally addressed in model organisms representing just a few clades. Recent efforts have been made to extend the characterization from differential expression to epigenetics in the less represented groups like butterflies, moths, mosquitoes, and lizards^100,116–119^. By combining a phylogenetic analysis of the *C. elegans* DCC with a comparative analysis of its genomic, transcriptomic, and epigenomic signatures, our work found that a new dosage compensation mechanism evolved in the presence of an existing one in the *Caenorhabditis* lineage.

The existing dosage compensation mechanism in the common ancestor of *O. tipulae*, *C. elegans*, and *P. pacificus* may be linked to H4K20me1, and may have constrained evolution towards an X-specific condensin in *Caenorhabditis* and *Pristionchus*. Furthermore, the observation that SMC proteins repeatedly evolved for dosage compensation in nematodes *and* mammals argues for a fundamental link between the chromosome-wide activity of SMC complexes and the need to regulate transcription across the entire X chromosome. A more exhaustive accounting of dosage compensation phenotypes in multiple species spanning *all* five clades would help delineate the history of sex chromosome dosage compensation in nematodes, and potentially uncover new strategies for chromosome wide gene regulation.

## AUTHOR CONTRIBUTIONS

A.A. and A.W. made the phylogenetic trees. A.A. and J.K. generated the Hi-C libraries. A.A. collected all samples, generated the ChIP-seq libraries, analyzed the sequencing data, performed nigon painting, re-scaffolded and lifted over the *P. pacificus* genome, ran the statistical analyses, and prepared the figures. A.A. and S.E. conceived the project and wrote the manuscript.

## ACKNOWLEDGEMENTS

We thank David Fitch and Karin Kiontke for their discussions on nematode phylogenetics. We thank Sophie Tintori (O), Michael Werner (P), Bogdan Sierebriennikov (P), Menyi Cao (S), and Carly Meyers (S) for their advice on working with *O. tipulae*, *P. pacificus* and *S. hermaphroditum*. We thank Christian Roedelsperger for discussions on *Pristionchus* phylogeny and genomes. We thank George Chung for discussions on *Diploscapter* genomes, and for providing the annotations for *D. coronatus* and *D. pachys*. We thank Andre Pires da Silva for providing the annotations for *A. rhodense*. The CEW1 strain was provided by the Caenorhabditis Genetics Center (CGC). The PS9179 and HG2587 strains were generously provided by Mengyi Cao. Sequencing and processing of the raw data was done by the Genomics Core at the Center for Genomics and Systems Biology at New York University. AA was supported in part by NIGMS Predoctoral Fellowship T32HD007520, and AA and SE in part by R35GM130311. AW was supported by NSF REU Site award number 2148706.

---

**Figure S1.**
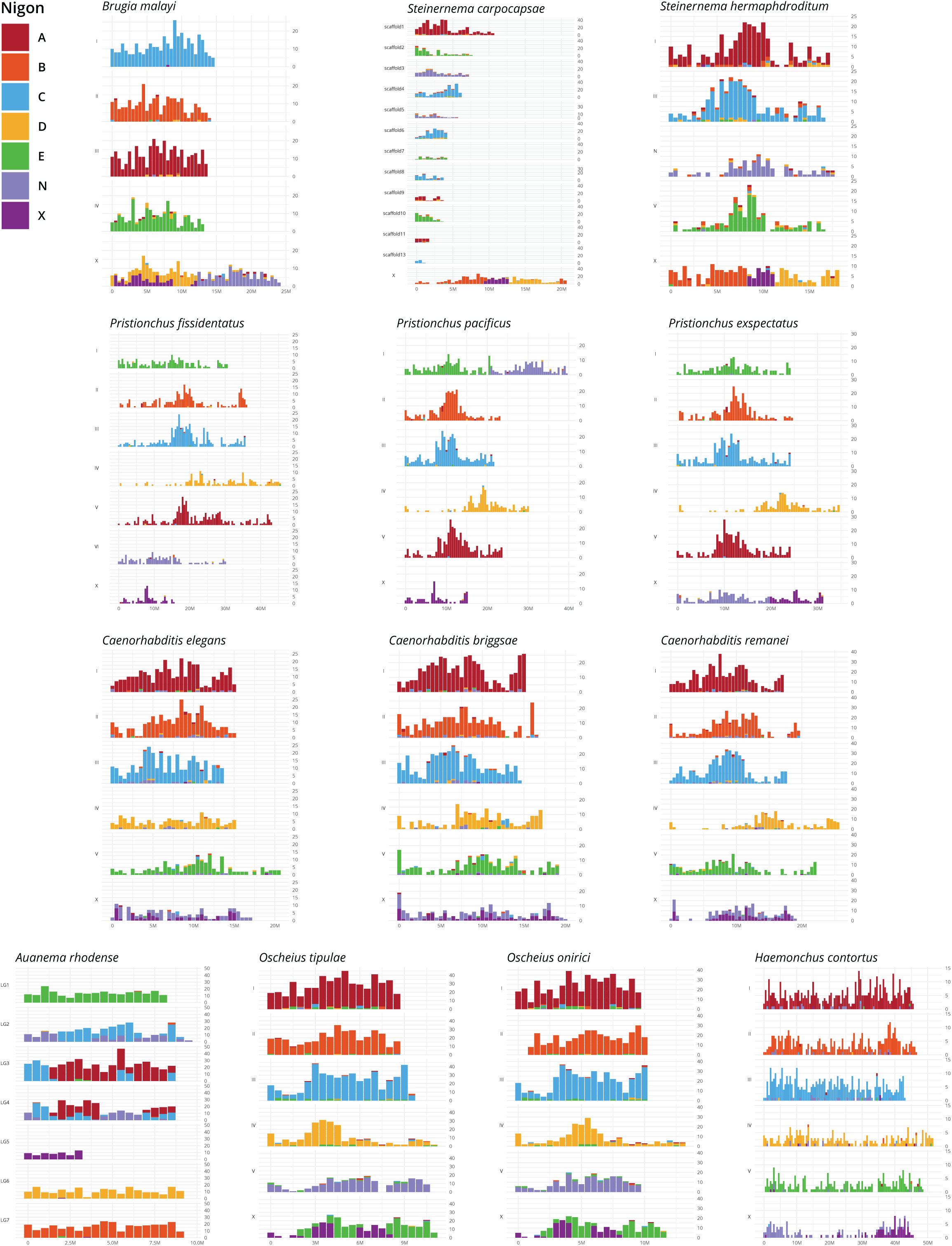
Nigon painted chromosomes of sampled nematodes. Chromosomes painted by ancestral nigon elements for species in Spirurina, Tylenchina, and Rhabditina support multiple independent autosome to sex chromosome fusions in nematodes, and significant rearrangements in nematode X chromosomes

**Figure S2.**
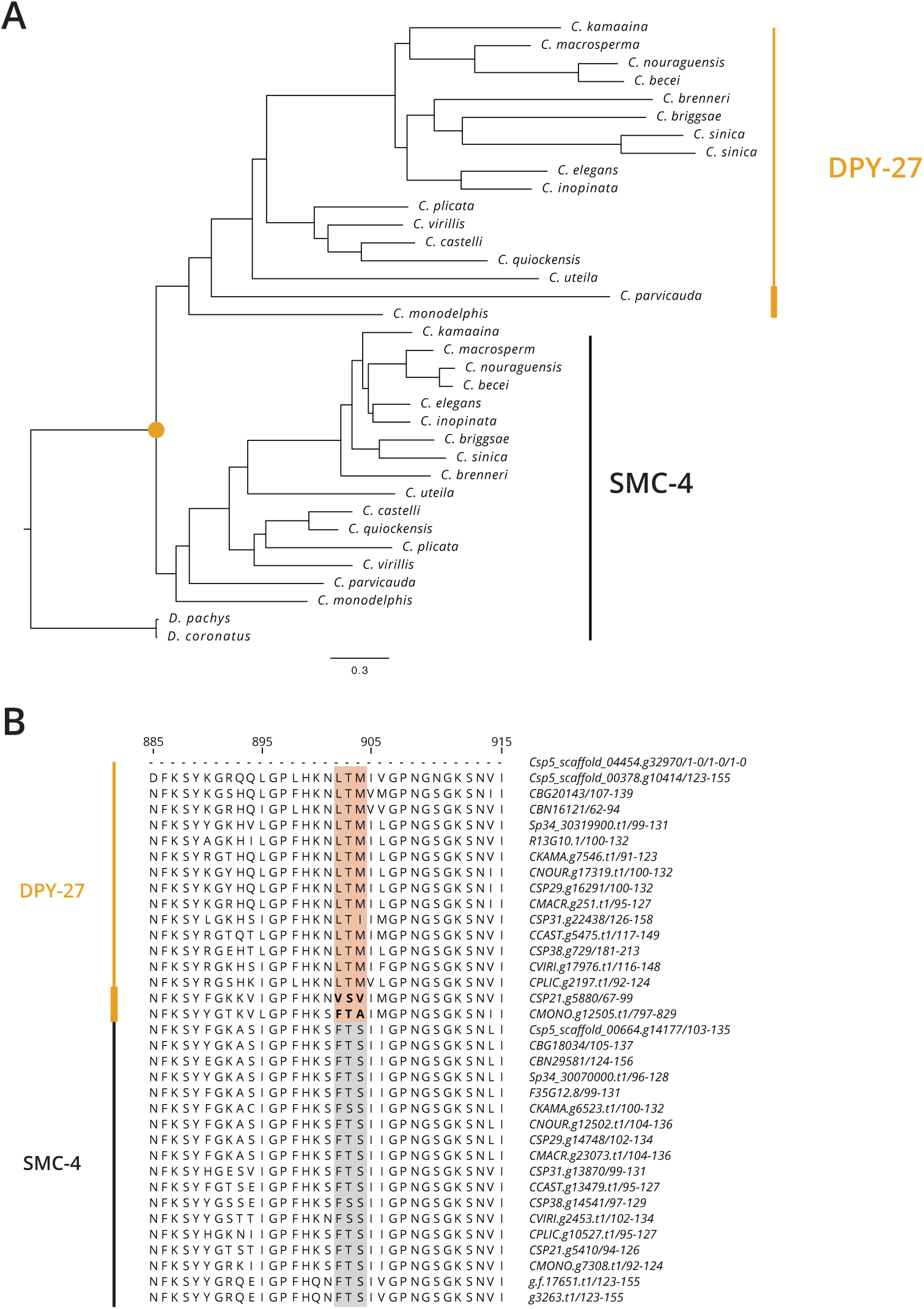
Maximum-likelihood tree of SMC-4 and DPY-27 in *Caenorhabditis*. (A) SMC-4 duplication occurred before the earliest known branch in *Caenorhabditis*, *C. monodelphis*, and after the split between *Caenorhabditis* and *Diploscapter*. (B) The LTM motif is exclusive to DPY-27 and is conserved after the *C. pavicuada* split.

**Figure S3.**
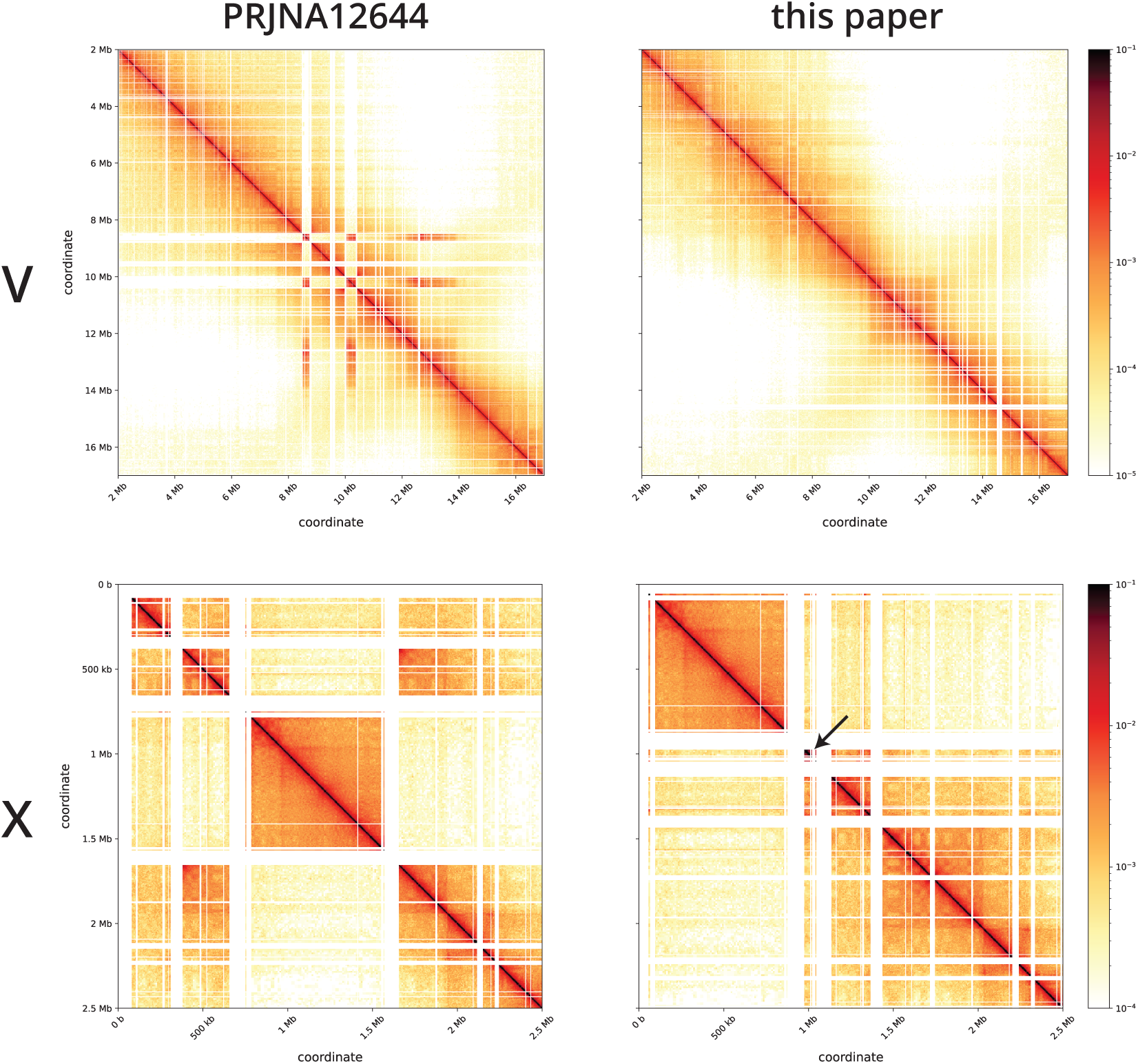
Fixed misjoins and inversions in *P. pacificus* genome. Misjoins and inversions (left panel) on chromosome V and X of the *P. pacificus* genome were fixed (right panel) with Juicebox Assembly Tools, and pbcontig517 was placed on the left arm of chromosome X (arrow) using the scaffolding tool YaHS.

**Figure S4.**
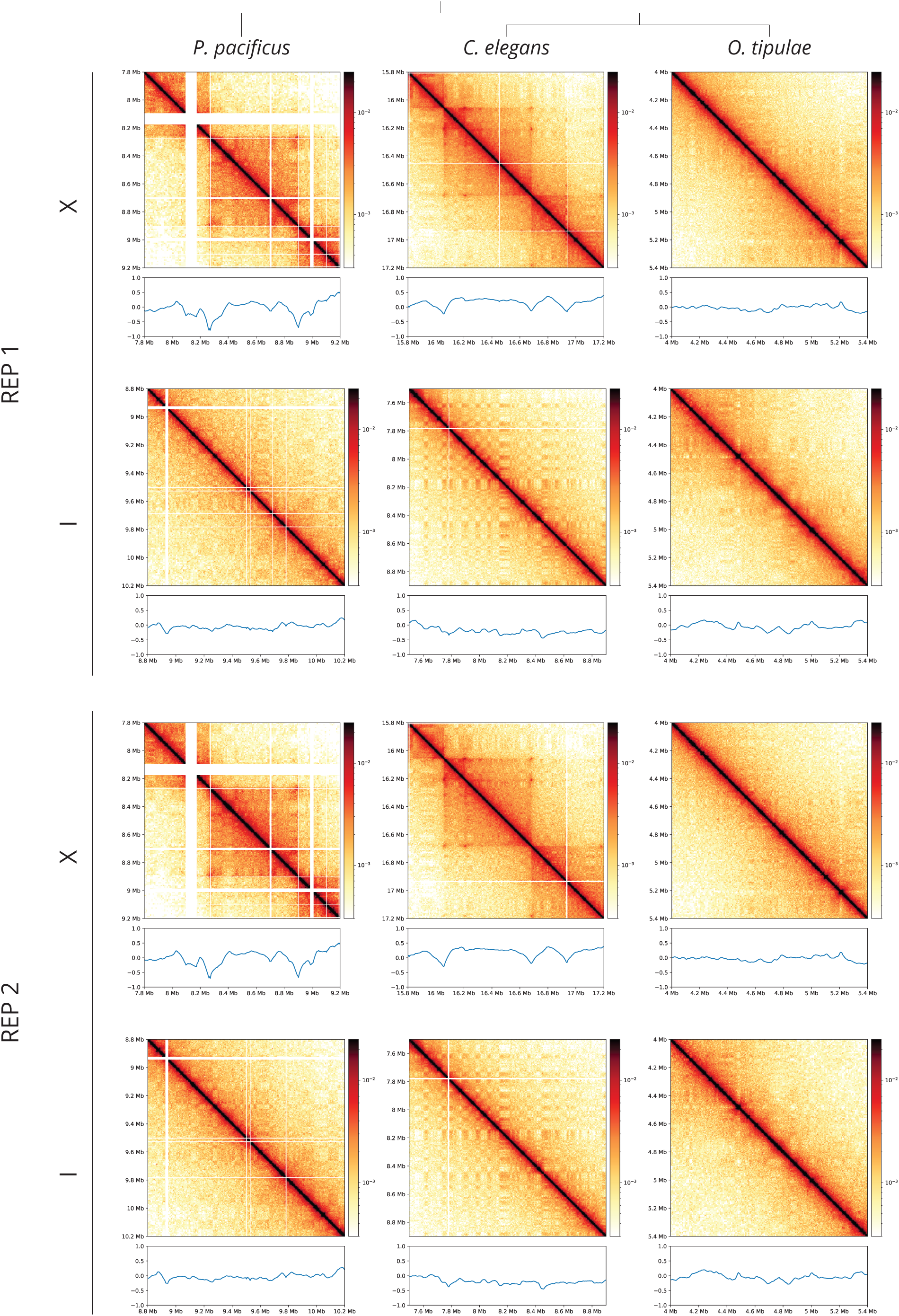
Replicates of Hi-C matrices and insulation scores. Replicates of Hi-C matrices (5 kb resolution) and insulation scores (150 kb window size) for data generated in this study exhibit reproducibility in displaying X chromosome specific TADs.

**Figure S5.**
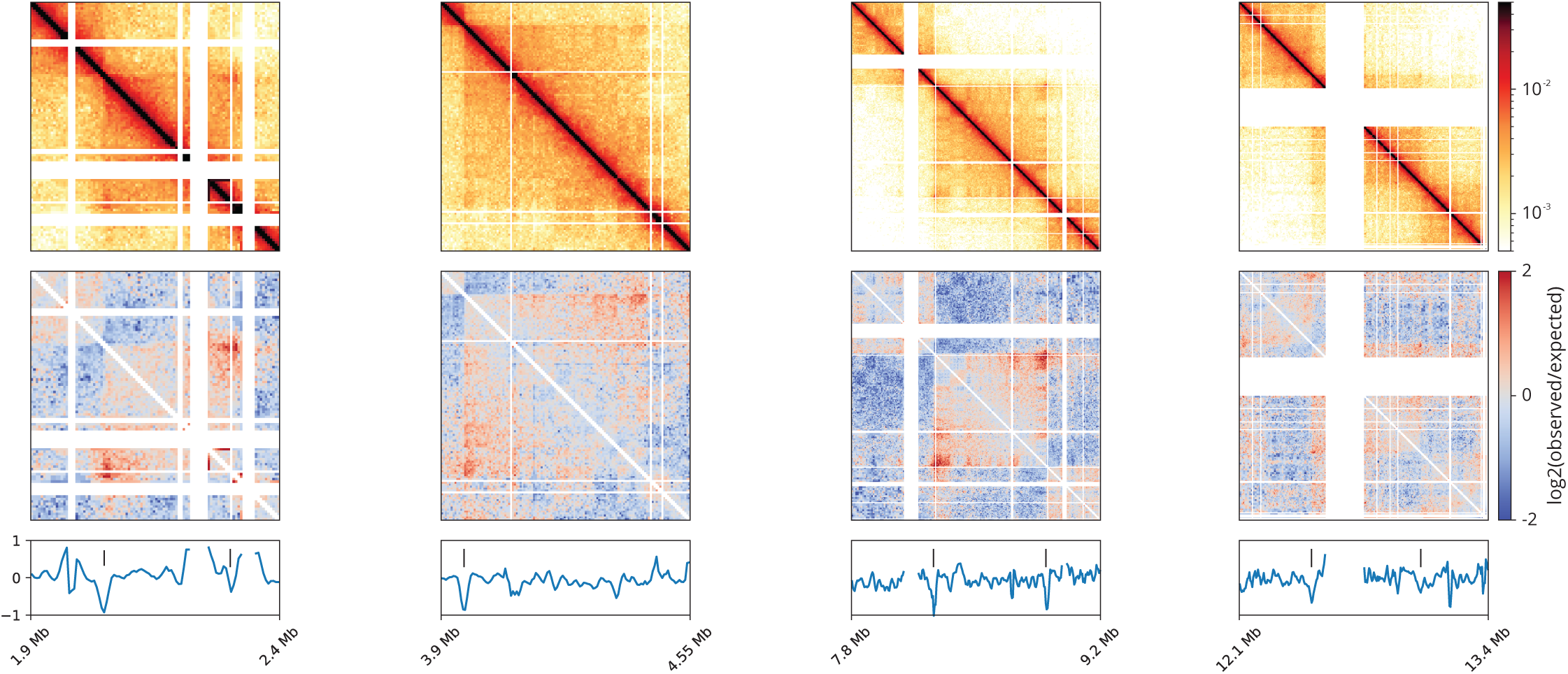
Hi-C matrices of the observed X chromosome TADs in *P. pacificus*. Hi-C matrices of each of the four observed TADs on the *P. pacificus* X chromosome (top panel) with log2(observed/expected) plots (middle panel). Dips in insulation score (bottom panel, 20 kb window size) represent TAD boundaries and are marked by ticks. Empty bins resulting in insulation score dips were not considered.

**Figure S6.**
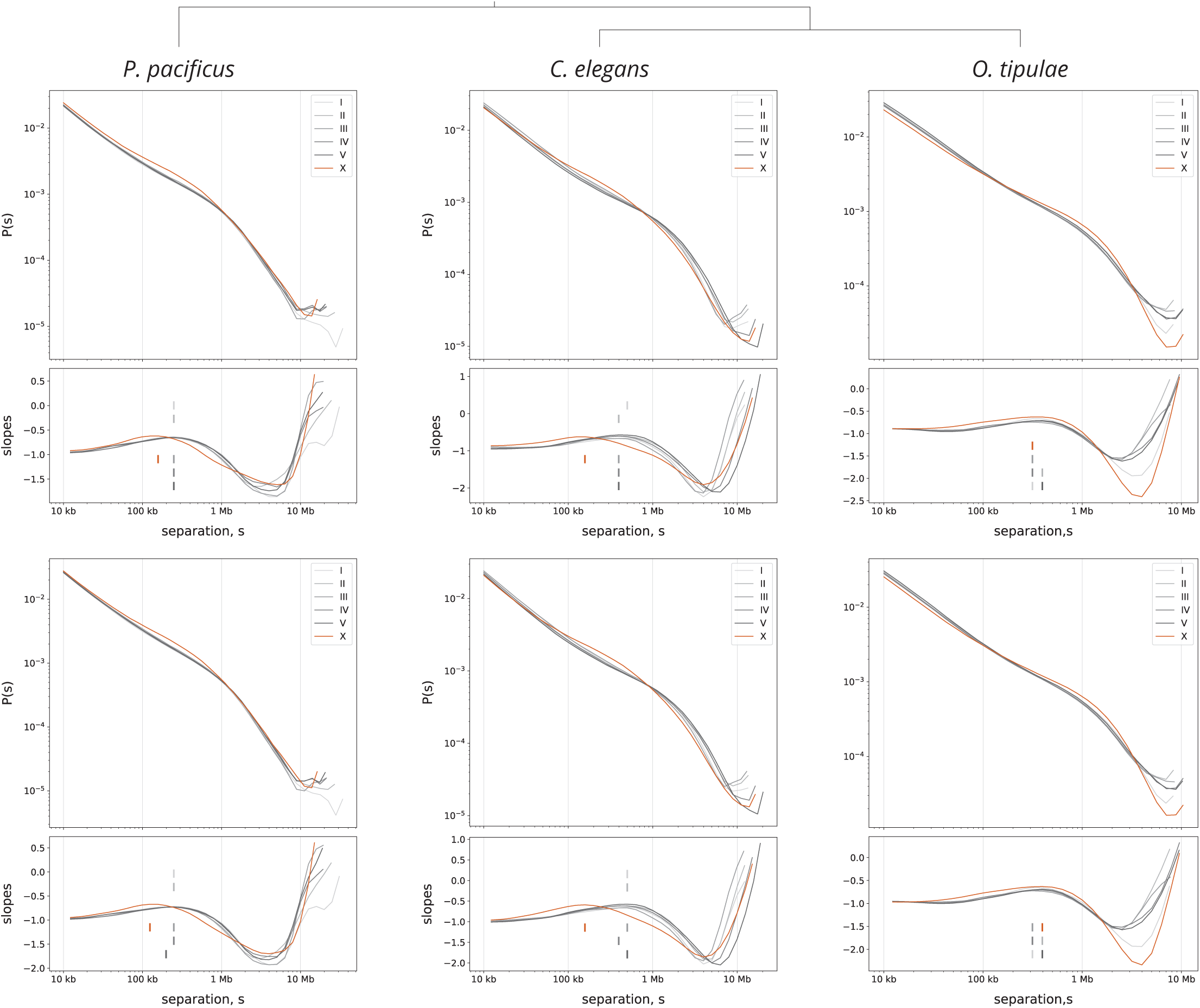
Replicates of distance decay curves. Replicates of P(s) curve and its derivative exhibit reproducibility of estimated mean loop size.

**Figure S7.**
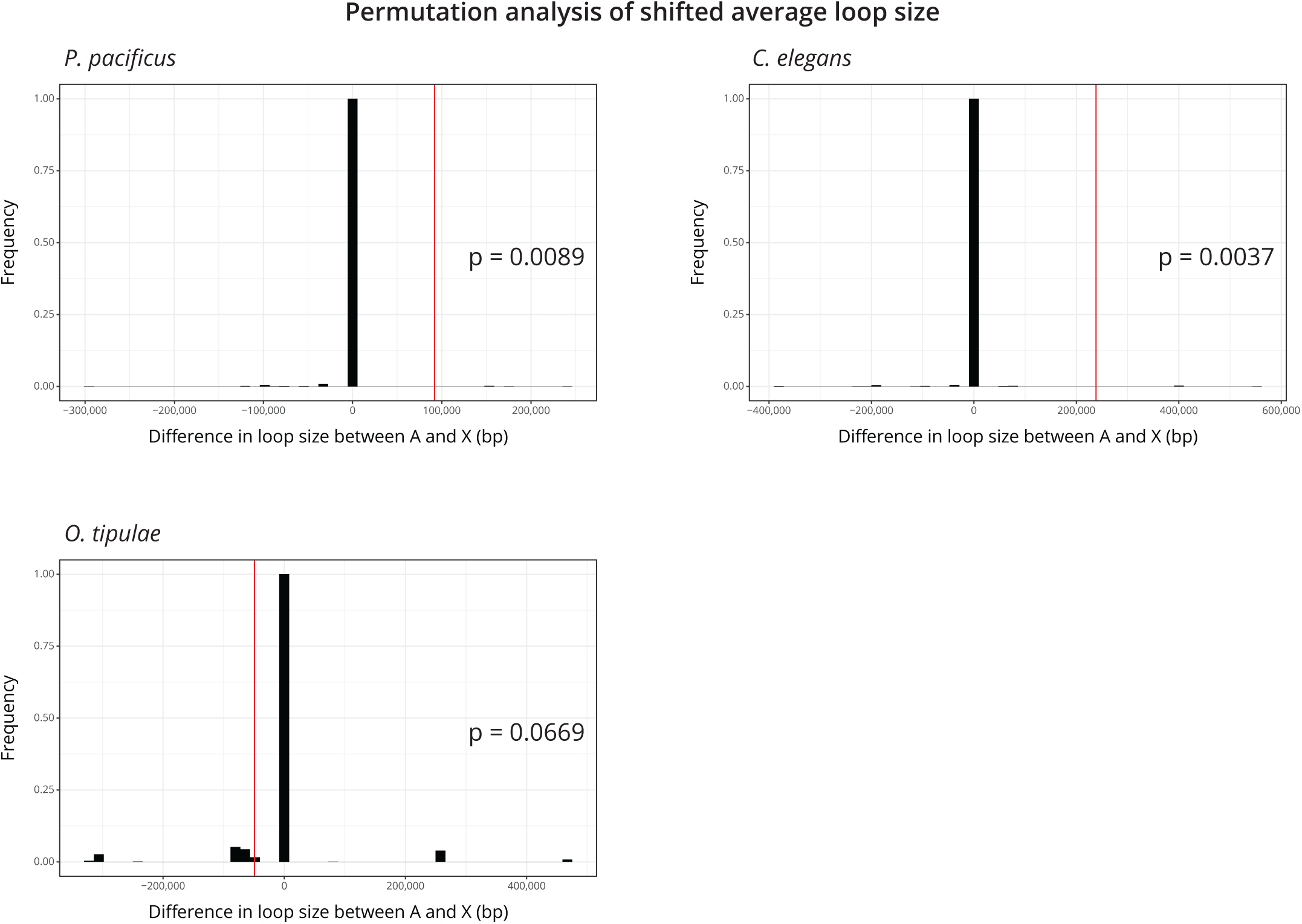
The observed difference in mean loop size between the autosomes and X chromosome are significantly different in *P. pacificus* and *C. elegans*. Permutations were performed by randomly permuting the slope values between the X chromosome and autosomes, extracting the corresponding separation in bp, and subtracting the X from the A as our test statistic. The distribution represents n=10000 permutations. Red line, observed difference; p, proportion of permutations that were more extreme than the observed difference.

**Figure S8.**
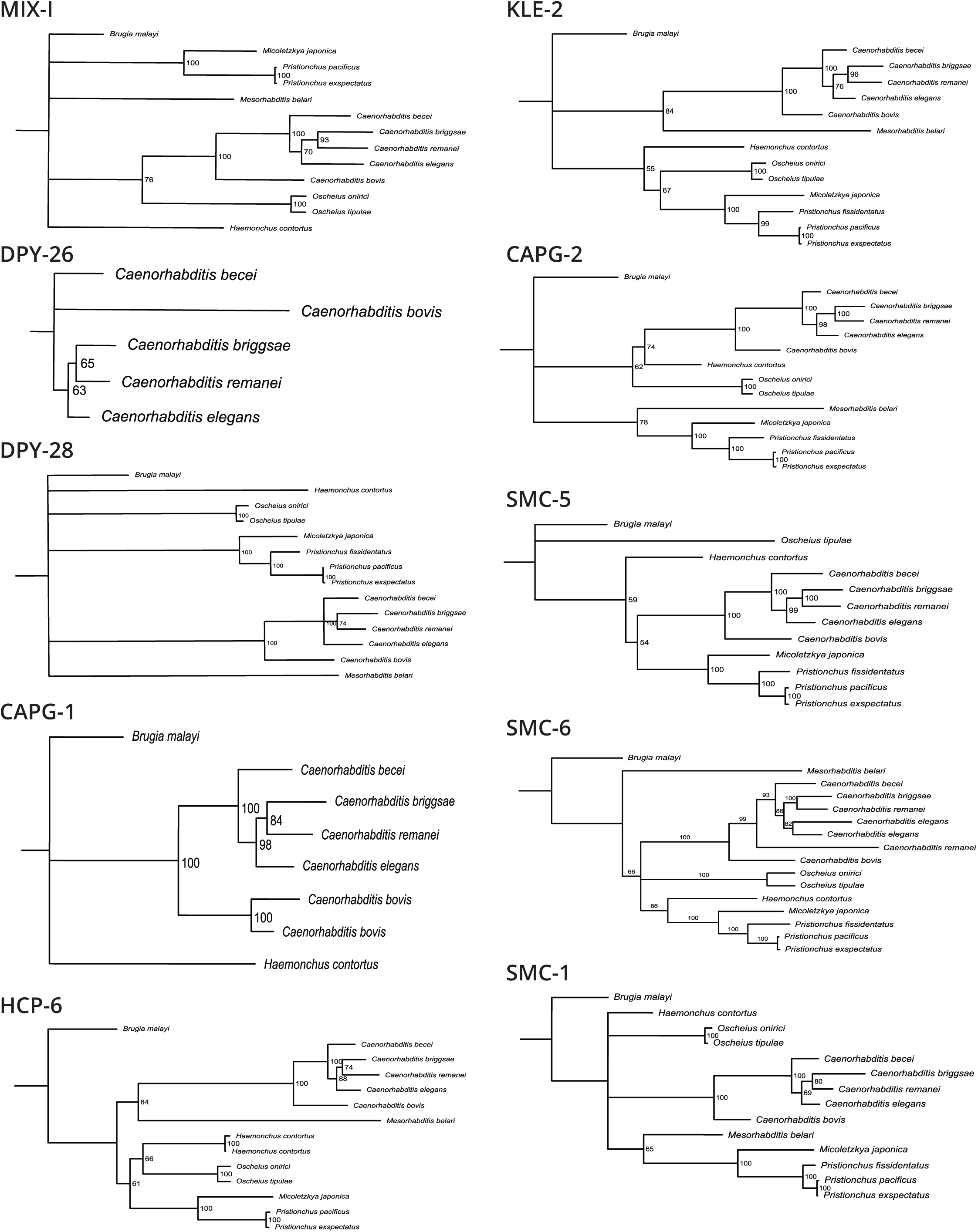
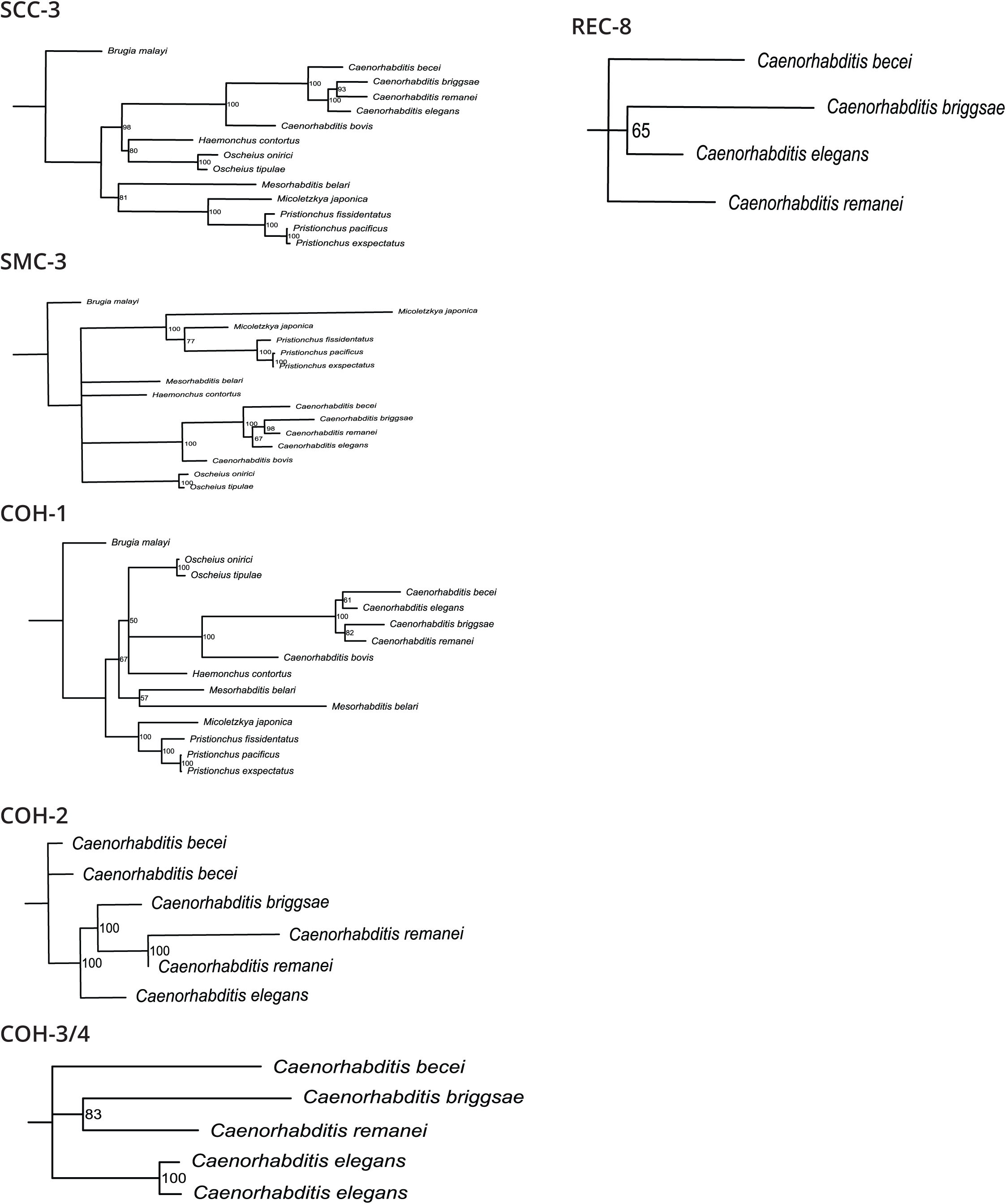
Maximum-likelihood trees of SMC proteins in nematodes. Maximum likelihood trees were generated for all condensin, cohesin, and SMC-5/6 subunits, which show no other SMC duplications in *P. pacificus*. See **supplemental file 1** for list of species used.

**Figure S9.**
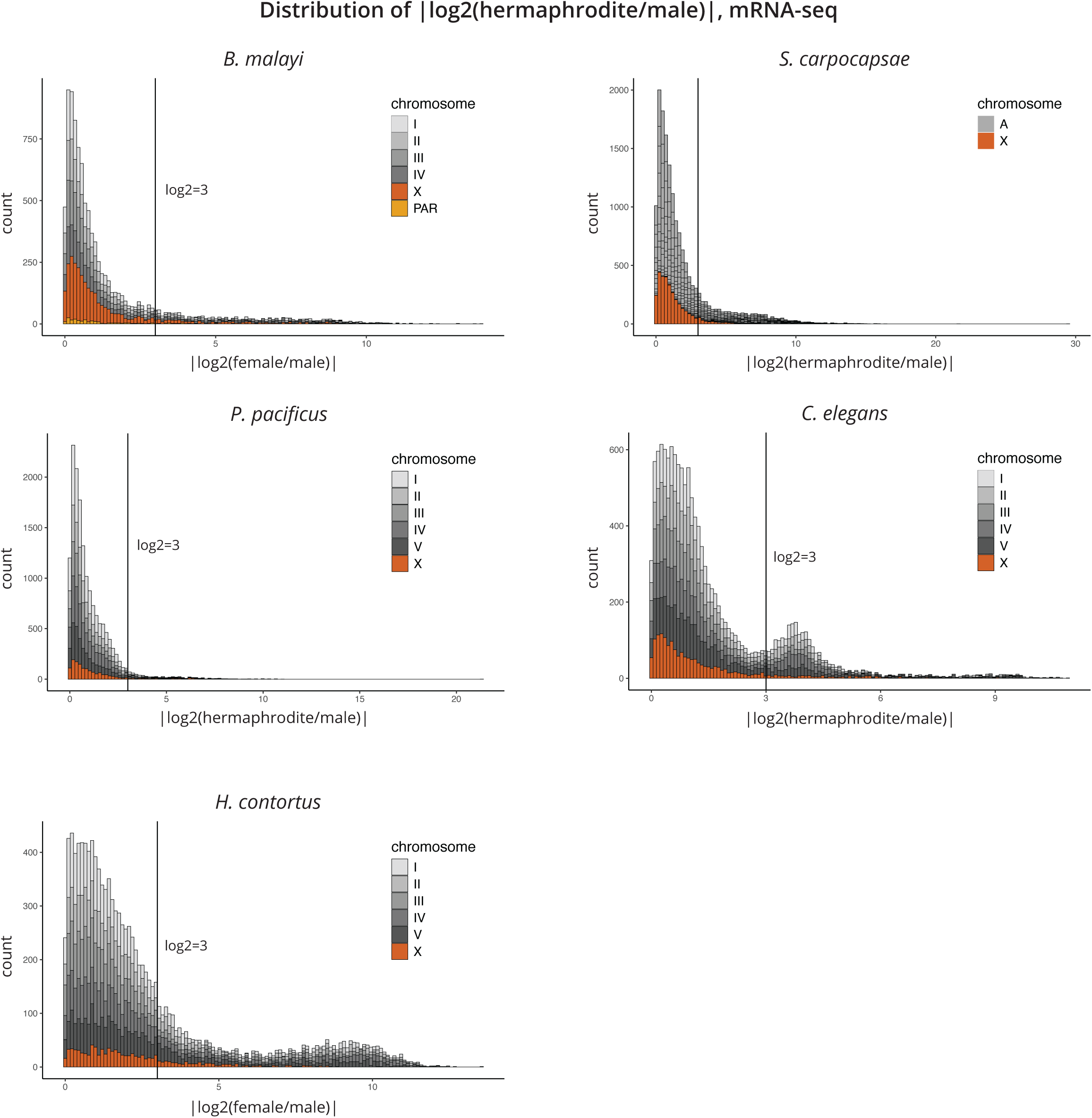
Distribution of |log2 fold change| in *B. malayi*, *S. carpocapsae*, *P. pacificus*, *C. elegans*, and *H. contortus*. The distribution of differentially expressed genes between sexes is split into soma (log2 < 3) and sex-biased, germline enriched genes (log2 > 3). The dearth of germline enriched genes in *P. pacificus* was addressed in Albritton et al. (2016)^84^.

**Figure S10.**
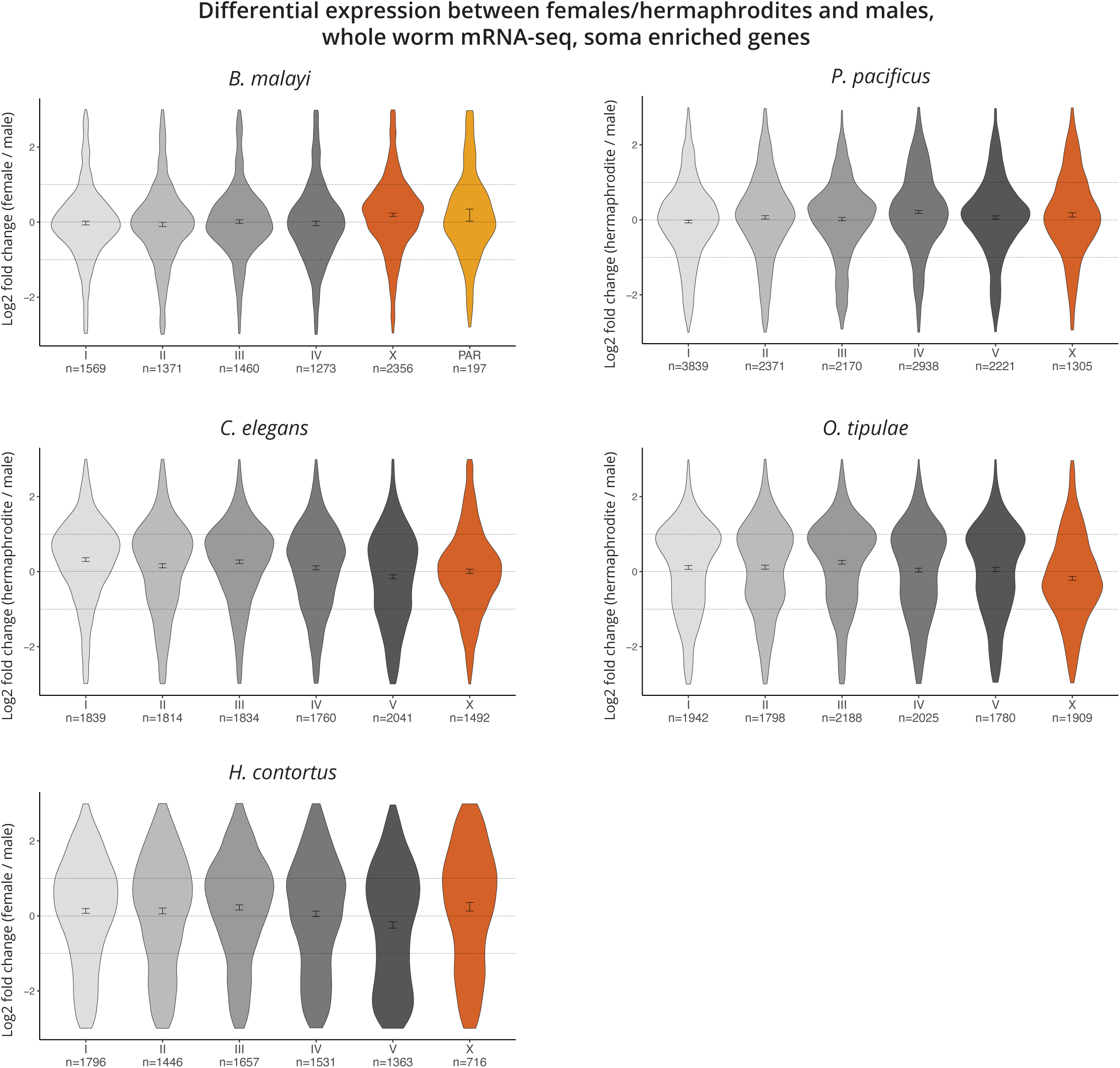
log2 fold change plots separated by chromosome. Log2 fold change between female (or hermaphrodite) and males of soma enriched expressed genes in *B. malayi*, *P. pacificus*, *C. elegans*, *O. tipulae*, and *H. contortus* show little to no difference between chromosomes. PAR, pseudoautosomal region.

**Figure S11.**
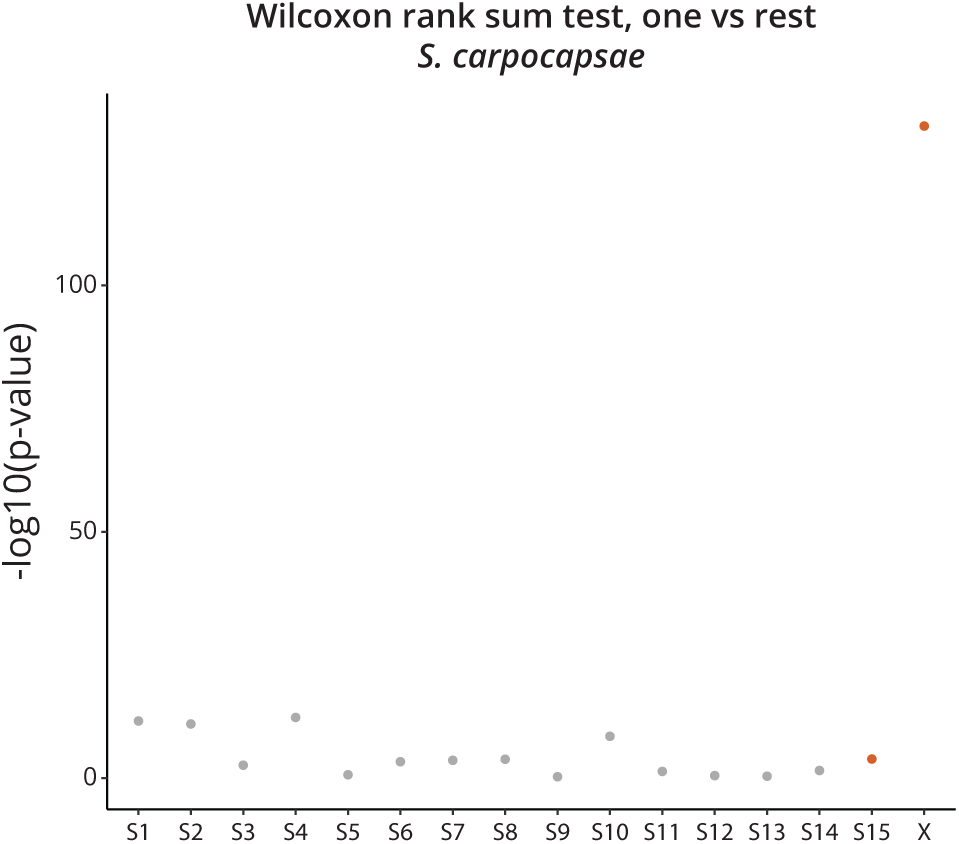
*S. carpocapsae* X chromosomes display significant differential expression when compared to autosomes. -log10(p-value) from Wilcoxon rank sum tests comparing the mean log2 fold change of a scaffold/chromosome to the mean of the rest (e.g., mean of X and mean of S1-14). Here, the differential expression observed between sexes on the X chromosome cannot be explained by the existing variation in autosomes between sexes.

**Figure S12.**
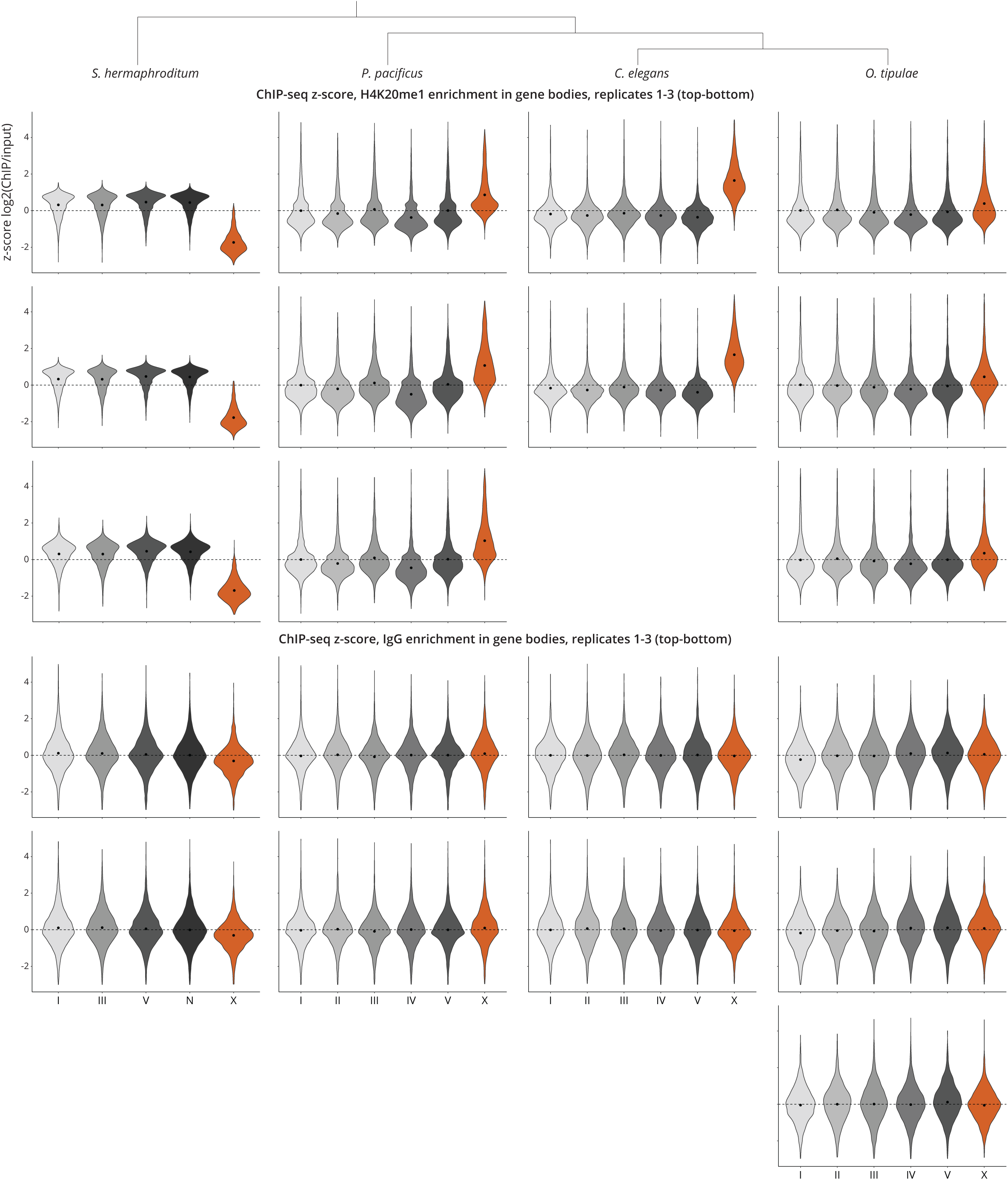
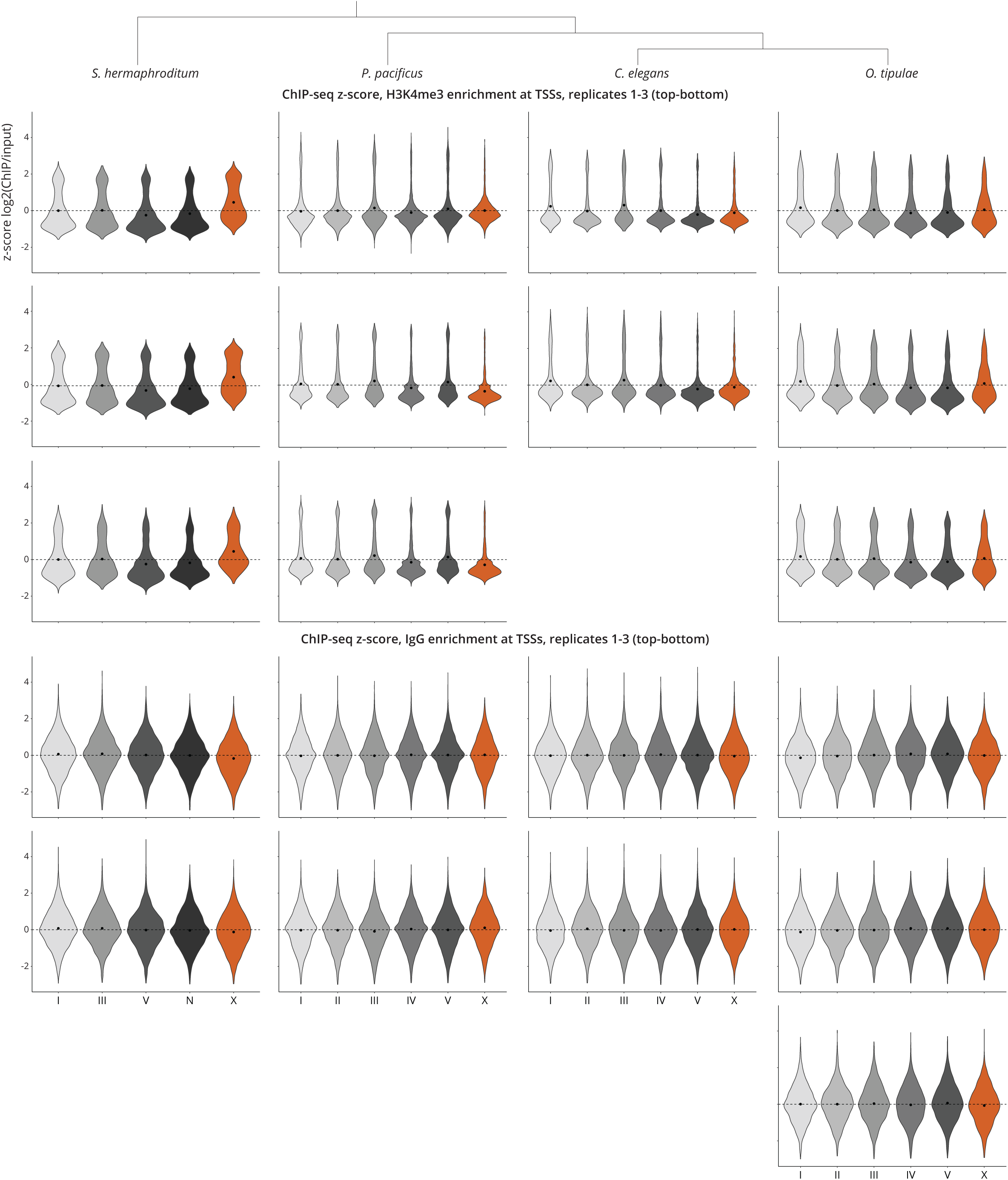
Replicates of enrichment of H4K20me1 at gene bodies and H3K4me3 at TSSs. Replicates of ChIP-seq z-score analysis exhibit reproducibility. IgG is shown as a negative control.

**Figure S13.**
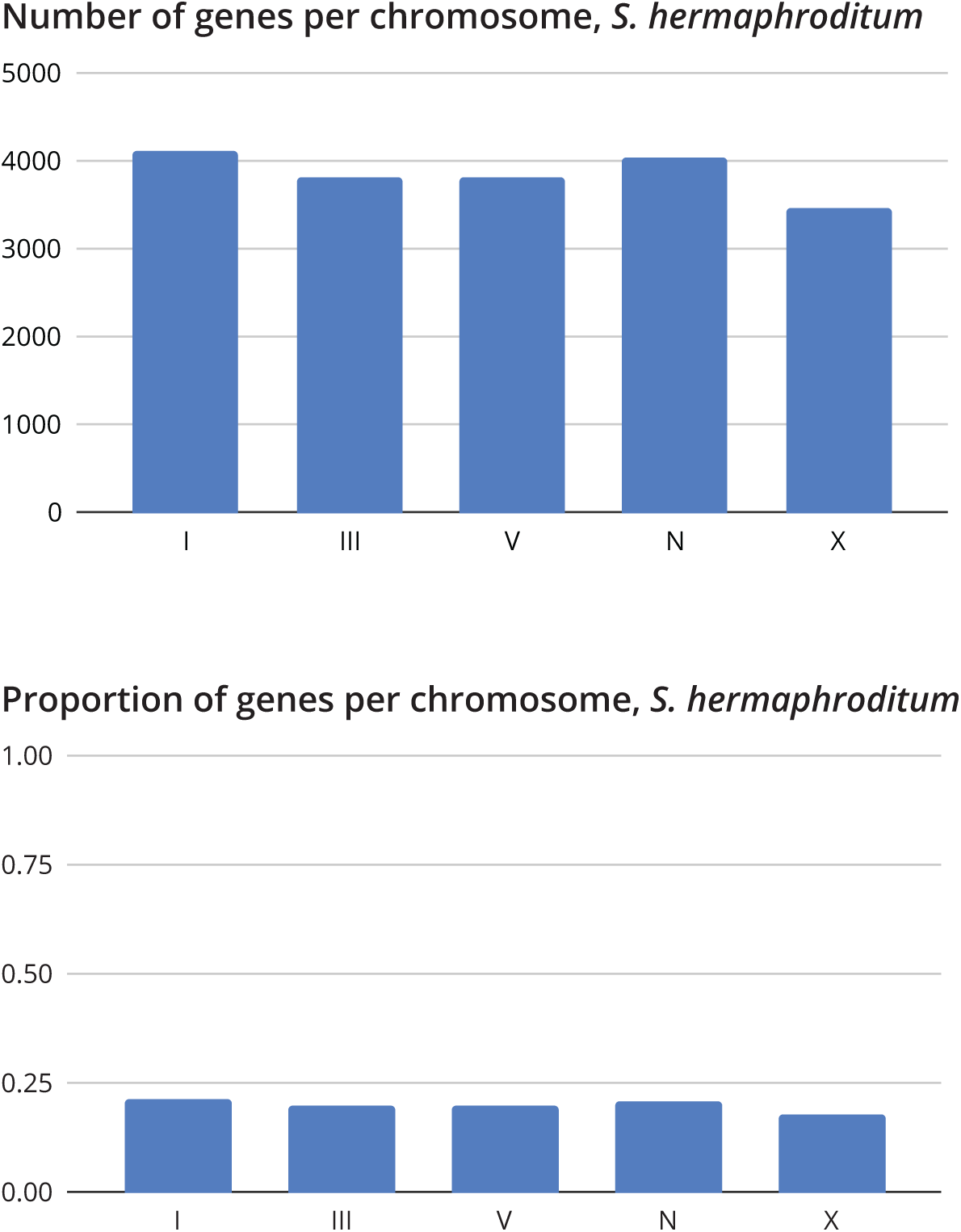
Distribution of genes on the *S. hermaphroditum* chromosomes. Genes per chromosome were plotted. Genes in *S. hermaphroditum* are distributed equally among its five chromosomes, and the distribution cannot explain the de-enrichment of H4K20me1 on the X chromosomes.

